# Nanoscale Curvature Regulates YAP/TAZ Nuclear Localization Through Nuclear Deformation and Rupture

**DOI:** 10.1101/2024.10.18.619165

**Authors:** Emmet A. Francis, Einollah Sarikhani, Vrund Patel, Dhivya Pushpa Meganathan, Leah Sadr, Zeinab Jahed, Padmini Rangamani

## Abstract

Nuclear translocation of the transcription regulatory proteins YAP and TAZ is a critical readout of cellular mechanotransduction. Recent experiments have demonstrated that cells on substrates with well-defined nanotopographies demonstrate mechanoadaptation through a multitude of effects - increased integrin endocytosis as a function of nanopillar curvature, increased local actin assembly on nanopillars but decreased global cytoskeletal stiffness, and enhanced nuclear deformation. How do cells respond to local nanotopographical cues and integrate their responses across multiple length scales? This question is addressed using a biophysical model that incorporates plasma membrane (PM) curvature-dependent endocytosis, PM curvature-sensitive actin assembly, and stretch-induced opening of nuclear pore complexes (NPCs) in the nuclear envelope (NE). This model recapitulates lower levels of global cytoskeletal assembly on nanopillar substrates, which can be partially compensated for by local actin assembly and NE indentation, leading to enhanced YAP/TAZ transport through stretched NPCs. Using cell shapes informed by electron micrographs and fluorescence images, the model predicts lamin A and F-actin localization around nanopillars, in good agreement with experimental measurements. Finally, simulations predict nuclear accumulation of YAP/TAZ following rupture of the NE and this is validated by experiments. Overall, this study indicates that nanotopography tunes mechanoadaptation through both positive and negative feedback on mechanotransduction.

**Table of Contents:** 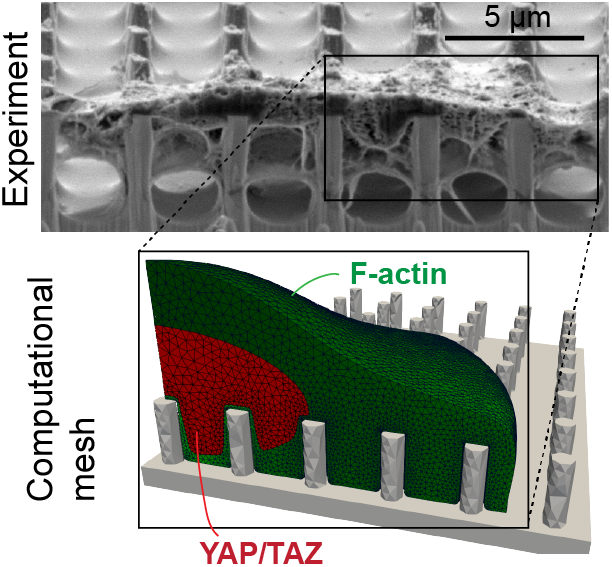

This study uses experiments and biophysical modeling to examine the response and adaptation of cells to nanoscale topography of surfaces. It is shown that cytoskeletal assembly and nuclear localization of transcription regulatory factors such as YAP/TAZ can be tuned by nanoscale membrane curvature and nuclear deformation and rupture due to substrate nanotopography.

## 1 Introduction

Mechanotransduction, the set of processes by which cells respond to mechanical cues by altering their signaling response, is critical for integration of chemical and physical stimuli in different physiological environments. Given the wide range of stiffnesses exhibited by tissues throughout the human body^[1]^ and changes in mechanical properties of extracellular matrix (ECM) associated with diseases such as cancer and atherosclerosis and with aging,^[2, 3, 4]^ mechanotransduction occupies a pivotal role in cell physiology. However, the manner in which a cell simultaneously responds and adapts to combined mechanical cues such as stiffness, membrane curvature, and other micro- and nano-topographical features remains unclear. Collectively, such cell responses are referred to as mechanoadaptation and are critical for cell migration through diverse extracellular geometries,^[5, 6]^ which are often composed of dense fibrillar networks.^[7, 8, 9]^

Engineered substrates with nano-scale features such as nanobars and nanopillars have proven to be a valuable experimental tool to probe cellular responses to nanotopography.^[10, 11, 12]^ Such substrates induce positive (inward) curvature in regions of the plasma membrane (PM), leading to changes in local processes such as endocytosis,^[13, 14, 12]^ integrin recruitment,^[6]^ and PM-endoplasmic reticulum contact sites,^[15]^ as well as whole-cell properties such as cell stiffness.^[14]^ Recent studies indicate that the assembly and remodeling of cytoskeleton-associated molecules is a function of underlying substrate curvature. For instance, cells that spread over nanobars or nanopillars formed adhesion complexes distinct from conventional focal adhesions.^[6, 16, 17]^

These differences in cell-ECM interactions and cytoskeletal assembly contribute to changes in nuclear localization of transcriptional regulatory proteins such as yes-associated protein (YAP) and the closely related protein, transcriptional coactivator with PDZ-binding motif (TAZ, also known as WWTR1). YAP and TAZ regulate gene expression by binding to transcriptional enhanced associate domains (TEADs) in the nucleus.^[18, 19]^ It was found that the extent of YAP nuclear translocation depended not only on substrate stiffness but also on substrate nanotopography.^[14]^ Specifically, cells grown on stiff substrates with nanopillars showed reduced cytoskeletal activity, similar to those grown on flat, soft substrates.^[14, 20]^ This was attributed to an increase in integrin endocytosis at inwardly curved regions of the PM, resulting in reduced cytoskeletal stiffness and, therefore, lower YAP nuclear abundance.^[14]^ However, nanotopography has also been indicated as a positive regulator of mechanotransduction, inducing local actin assembly in a curvature-dependent manner.^[17, 16]^ Furthermore, nanotopography may influence the nuclear transport of mechanosensitive factors through stretch^[21]^ and even rupture^[22]^ of the nuclear envelope (NE). These opposing effects of nanotopography on signaling events make it unclear to what extent mechanotransduction is positively or negatively regulated by nanoscale curvature in different contexts. Here, we use a combination of modeling and experiments to investigate how cells integrate nanotopographical cues to generate mechanotransduction responses.

Computational models have previously been used to predict the effects of substrate stiffness, substrate dimensionality, and ligand micropatterns on cytoskeletal activation and YAP/TAZ nuclear translocation in well-mixed and spatial frameworks.^[23, 24, 25, 26, 27]^ While these models shed light on the complex interplay between cell geometry and substrate stiffness in cell mechanotransduction, the role of substrate topography and its impact on YAP/TAZ nuclear translocation remain unexplored. Recent development of software to simulate reaction-transport networks in realistic cell geometries^[27, 28]^ enables us to conduct detailed simulations examining the effects of nanoscale substrate features on mechanotransduction signaling. We first generate experimentally informed model geometries corresponding to cells on different nanopillar substrates and run a series of simulations to calibrate our model to previous measurements of nuclear YAP as a function of nanopillar size and spacing.^[14]^. We then extend this model to describe nuclear transport of YAP/TAZ following nuclear deformation on nanopillars, relating this to direct measurements of nuclear indentation in U2OS cells. Finally, we use our model to predict the effect of NE rupture on YAP/TAZ signaling and then validate these predictions against experimental measurements, establishing that nuclear rupture can enhance YAP/TAZ nuclear accumulation on nanopillar substrates.

## 2 Results

### 2.1 Model summary

Building on previous models in the literature,^[24, 23]^ we model YAP/TAZ nuclear translocation as a function of nanotopography. Here, we present a brief summary of the main events included in our baseline model on a flat substrate. We will later introduce additional curvature- and deformation-dependent reactions at the PM and NE as needed.

Initially, tyrosine residues of focal adhesion kinase (FAK) are phosphorylated downstream of integrin binding in a substrate-stiffness-dependent manner.^[29, 30]^ FAK activates the small GTPase, Ras homolog family member A (RhoA) and Rho-associated protein kinase (ROCK), leading to actin polymerization and changes in myosin activity (Figure 1A). The formation of stress fibers is assumed to activate and free YAP/TAZ within the cytosol in a manner mediated by angiomotins and modulation of large tumor suppressor (LATS) kinases.^[31, 32]^ Since we focus on a single cell, we neglect other changes in LATS-mediated phosphorylation of YAP/TAZ due to formation of cell-cell contacts or other activation of Hippo signaling. Free cytosolic YAP/TAZ translocates into the nucleus through NPCs at a rate that depends on the extent of cytoskeletal assembly.^[33]^ In particular, it is assumed that lamin A is increasingly dephosphorylated as a function of cytosolic stiffness (a function of F-actin), causing it to be retained at the NE^[34]^ and interact with linker of nucleoskeleton and cytoskeleton (LINC) complexes to transmit force to NPCs^[35]^

**Figure 1.**
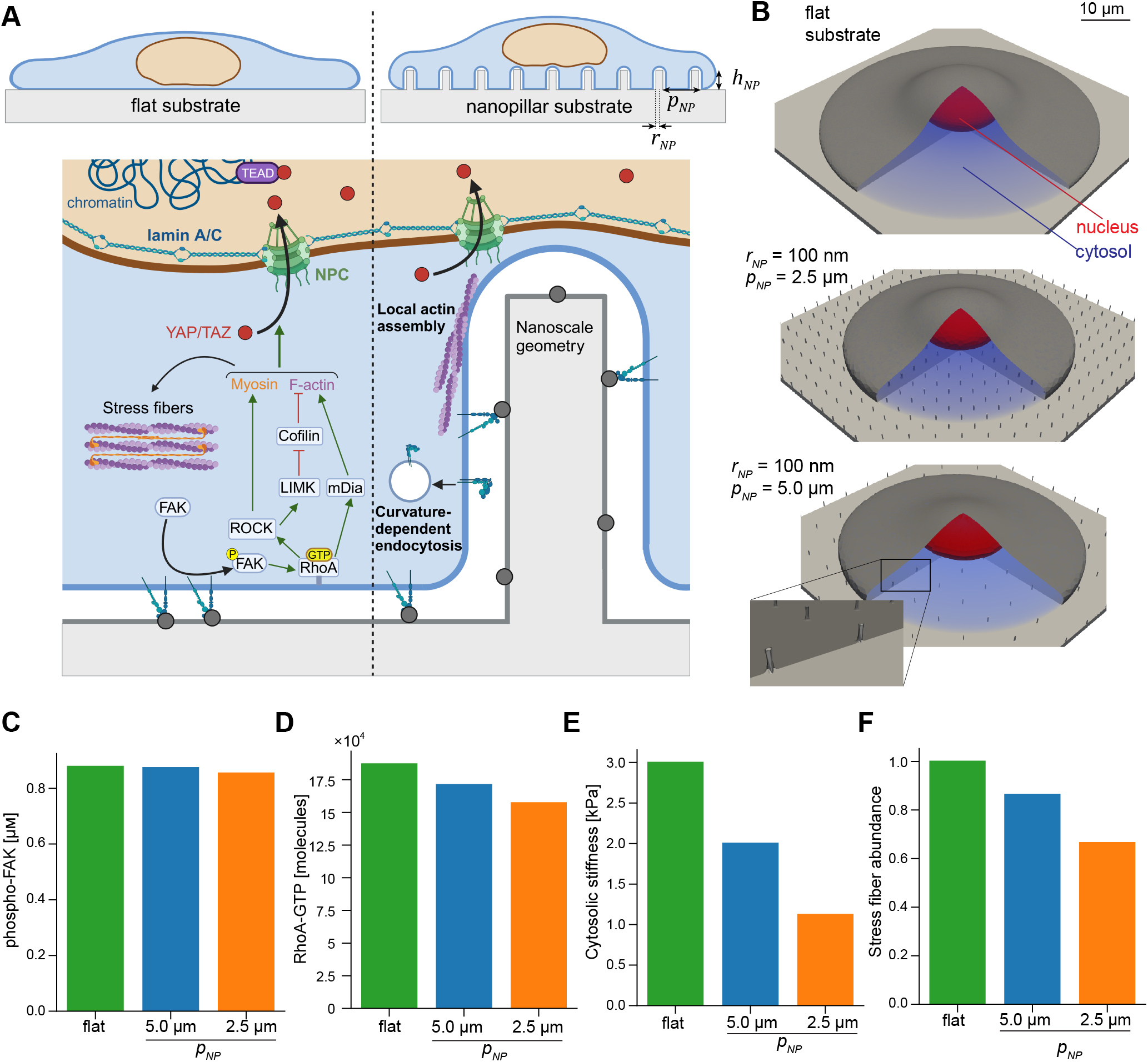
Substrate nanotopography alters activation of enzymes at the plasma membrane and downstream cytoskeletal activation. A) Schematic summarizing the main biochemical events leading to changes in nuclear YAP/TAZ in our model, in the case of a flat substrate (left), and additional signaling events introduced on nanopillar substrates (right). The arrangement of nanopillars is dictated by their height (*h*_*NP*_), pitch (*p*_*NP*_), and radius (*r*_*NP*_), which then affect signaling at the PM in a curvature-dependent manner, by enhancing N-WASP-mediated actin assembly and integrin endocytosis. B) Cell geometries on a flat substrate (upper) and nanopillar substrates with *p*_*NP*_ = 2.5 µm (middle) or 5.0 µm (lower), both with *r*_*NP*_ = 100 nm and *h*_*NP*_ = 1.0 µm. C-F) Phosphorylated FAK concentration, RhoA-GTP molecules at the PM, cytosolic stiffness (proportional to [*FActin*]^2.6^, as defined in previous work,^[24]^) and relative stress fiber abundance in the cytosol (proportional to [*FActin*][*Myo*_*A*_]) for all 3 geometries in panel B. Quantities were spatially averaged or integrated over the entire relevant compartment and assessed after reaching steady-state (*t* = 10,000 s) for *H*_0_ = 5 µm^*−*1^ and *k*_*N*-*W ASP*_ = 0.01 µm s^*−*1^.

These events are modeled deterministically using a system of mixed-dimensional partial differential equations governing the reaction and diffusion of each species (Section S2). We simulated these reaction-diffusion equations using our recently developed software package, Spatial Modeling Algorithms for Reaction and Transport (SMART).^[27, 28]^ The system is initialized to a state with minimal cytoskeletal activation and low levels of YAP/TAZ in the nucleus. Details on model implementation in SMART, as well as mesh generation using the mesh generator Gmsh,^[36]^ are provided in Methods and Supplementary Information.

### 2.2 Regulation of enzyme activation and actin polymerization by plasma membrane curvature

We first considered the effects of nanopillar size and spacing on YAP/TAZ signaling. Drawing from recent data showing inhibition of cytoskeletal activity on nanopillar substrates,^[14]^ we adapted our computational model to predict the effects of PM curvature-dependent integrin endocytosis (Figure 1A). In regions of high inward membrane curvature, endocytosis lowers the integrin density according to the following empirical relationship between bound integrin density, *ρ*_*I,bound*_, and mean curvature of the PM, *H*_*PM*_, in the membrane in contact with the substrate (Γ_*substrate*_):

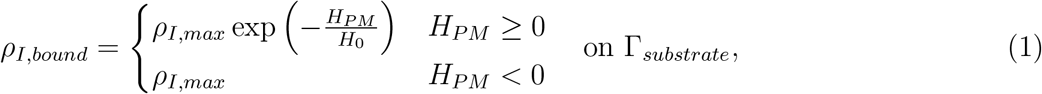

where *ρ*_*I,max*_ is the maximum density of bound integrins and *H*_0_ is a parameter controlling the degree of curvature sensitivity. *H*_*PM*_ was computed via Equation S36, as described in Section S4. Similar exponential relationships between reaction rates and membrane curvature have been used in previous models.^[16, 37]^

In line with previous iterations of the model,^[23]^ the FAK phosphorylation rate then depends on the local density of integrin-ligand complexes and on substrate stiffness (Young’s modulus), *E*_*mod*_:

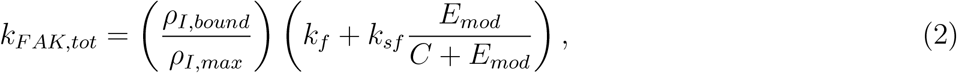

where *k*_*f*_ and *k*_*sf*_ are the stiffness-independent and stiffness-dependent FAK phosphorylation rates, respectively, and *C* is the stiffness sensitivity.

Separate studies have shown that F-actin assembles at highly curved regions of PM^[6, 16, 17]^ This has been attributed in part to formin-binding protein 17 (FBP17)-dependent activation of Neuronal Wiskott– Aldrich syndrome protein (N-WASP), leading to actin polymerization around nano-curved structures.^[16, 17]^ Accordingly, we introduce curvature-dependent actin polymerization at the PM in contact with the substrate:

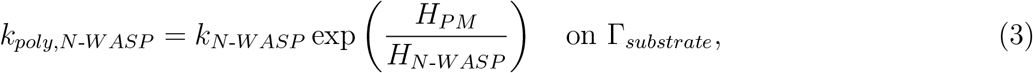

where *k*_*N*-*W ASP*_ is the rate of actin polymerization on flat PM and *H*_*N*-*W ASP*_ is the characteristic curvature governing the exponential relationship. Based on experimental measurements showing that N-WASP activation preferentially occurs on nanobars less than 500 nm in width, we fix *H*_*N*-*W ASP*_ at 2 µm^*−*1^ (the mean curvature of the curved sides of 500 nm nanobars). In what follows, we treat *k*_*N*-*W ASP*_ and *H*_0_ as free parameters; all other parameters were fixed to their previously estimated values,^[24]^ as reported in Table S4.

We conducted simulations using cell geometries that matched U2OS cell and nuclear volume and the contact areas measured for several of the nanopillar substrates tested by Li et al.^[14]^ These simulations showed that FAK phosphorylation was slightly lower on nanopillar substrates compared to flat substrates with the same stiffness (Figure 1C). Because the total contact area of the PM on nanopillar substrates is much greater than the projected area, the effect on FAK phosphorylation was relatively weak. However, the effect of curvature sensitivity was amplified for downstream RhoA activation (Figure 1D) due to the lower total PM surface area available for reaction in cells spread on nanopillar substrates (Table S1). Consequently, cytoskeletal activation was also appreciably lower, as indicated by the relative density of stress fibers and overall cytosolic stiffness, each defined as functions of activated myosin and/or F-actin (Figure 1E-F).

We observed that model cells on nanopillars exhibited enhanced local actin assembly as prescribed by Equation (3) (Figure 2A, Movies 1,2,3). However, because F-actin diffuses very slowly, this did not have a significant impact on global cytoskeletal assembly and only had a very small effect on nuclear YAP/TAZ (Figure S1). Overall, we observed that increases in nuclear YAP/TAZ correlated well with global cytoskeletal activation (Figure 2B-E, Movies 4,5,6). Smaller nanopillar spacing (higher nanopillar density) was associated with greater inhibition in cytoskeletal and YAP/TAZ activity. Over the values tested, *H*_0_ = 5 µm^*−*1^ and *k*_*W ASP*_ = 0.01 µm s^*−*1^ yielded the best match with experiments while still exhibiting local actin accumulation around nanopillars (Figure 2F, Figure S1). The best-fit value for *H*_0_ agrees well with the finding that clathrin-mediated endocytosis preferentially occurs on nanopillars less than 200 nm in radius,^[13]^ which induce a mean PM curvature of about 2.5 µm^*−*1^. Overall, this model predicts that the typical response of a cell to stiff ECM surfaces is inhibited on substrates with nanotopography.

**Figure 2.**
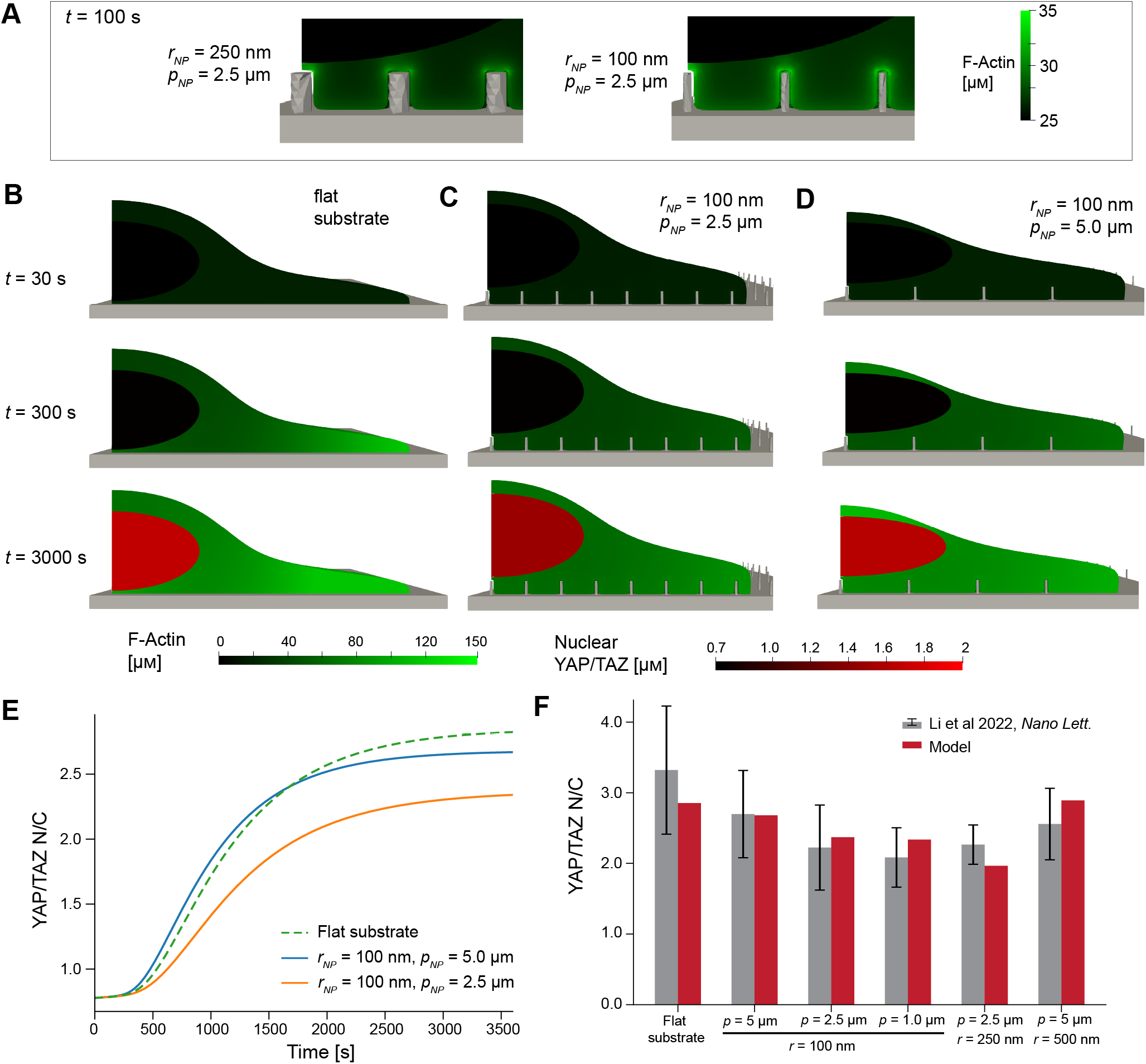
Simulations of nuclear translocation of YAP/TAZ on nanopillar substrates agree with experimental measurements. A) Magnified snapshot of the F-actin distribution around 250 nm or 100 nm nanopillars or above the flat substrate at *t* = 100 s. B-D) Spatial distributions of cytosolic F-actin (green) and nuclear YAP/TAZ (red) at 3 different time points for model cells on a flat substrate (B) or nanopillar substrates with *p*_*NP*_ = 2.5 µm (C) or 5.0 µm (D), both with *h*_*NP*_ = 1 µm and *r*_*NP*_ = 100 nm. Cell geometries were constrained by U2OS cell and nuclear volume and the average contact area measured previously on each substrate^[14]^ E) YAP/TAZ N/C dynamics for the three cases shown in B-D. F) Steady-state YAP/TAZ N/C for several nanopillar substrates tested in past experiments,^[14]^ compared to those predicted by our best-fit model where *H*_0_ = 5µm^*−*1^ and *k*_*W ASP*_ = 0.01 µm s^*−*1^. Experimental data is reported as mean ± standard deviation from Figure S10 in Li et al 2021.^[14]^ For these experimental measurements in gray, from left to right, *N* = 396, 38, 89, 59, 58, and 48.

### 2.3 Nanopillar-induced nuclear deformation leads to increases in the stretch and curvature of the nuclear envelope

Our previous studies showed that nuclear size and shape influence YAP/TAZ nuclear-to-cytosolic ratio (N/C) independently of global cell shape,^[24]^ so we next turned our attention to changes in nuclear morphology on nanopillar substrates. As previous studies have measured large deformations of the nucleus on nanopillar substrates,^[21, 22]^ we measured nuclear indentation and NE curvature in spread U2OS cells on nanopillar substrates. Cells were seeded on nanopillar substrates with base radius 0.585 ± 0.195 µm, center-to-center pitch 3.50 ± .08 µm, height 3.33 ± 0.21 µm, and tip radius 0.200 ± 0.0277 µm (mean ± standard deviation). U2OS nuclei were dramatically deformed, exhibiting indentations up to 3.5 µm in certain cases (Figure 3A-B). The curvature induced by such deformations ranged from 2.5-4.5 µm^*−*1^ (Figure 3B). We note that these nanopillars were much taller than those considered in the previous section (*>* 3 µm vs. 1 µm) in which we neglected nuclear indentation.

**Figure 3.**
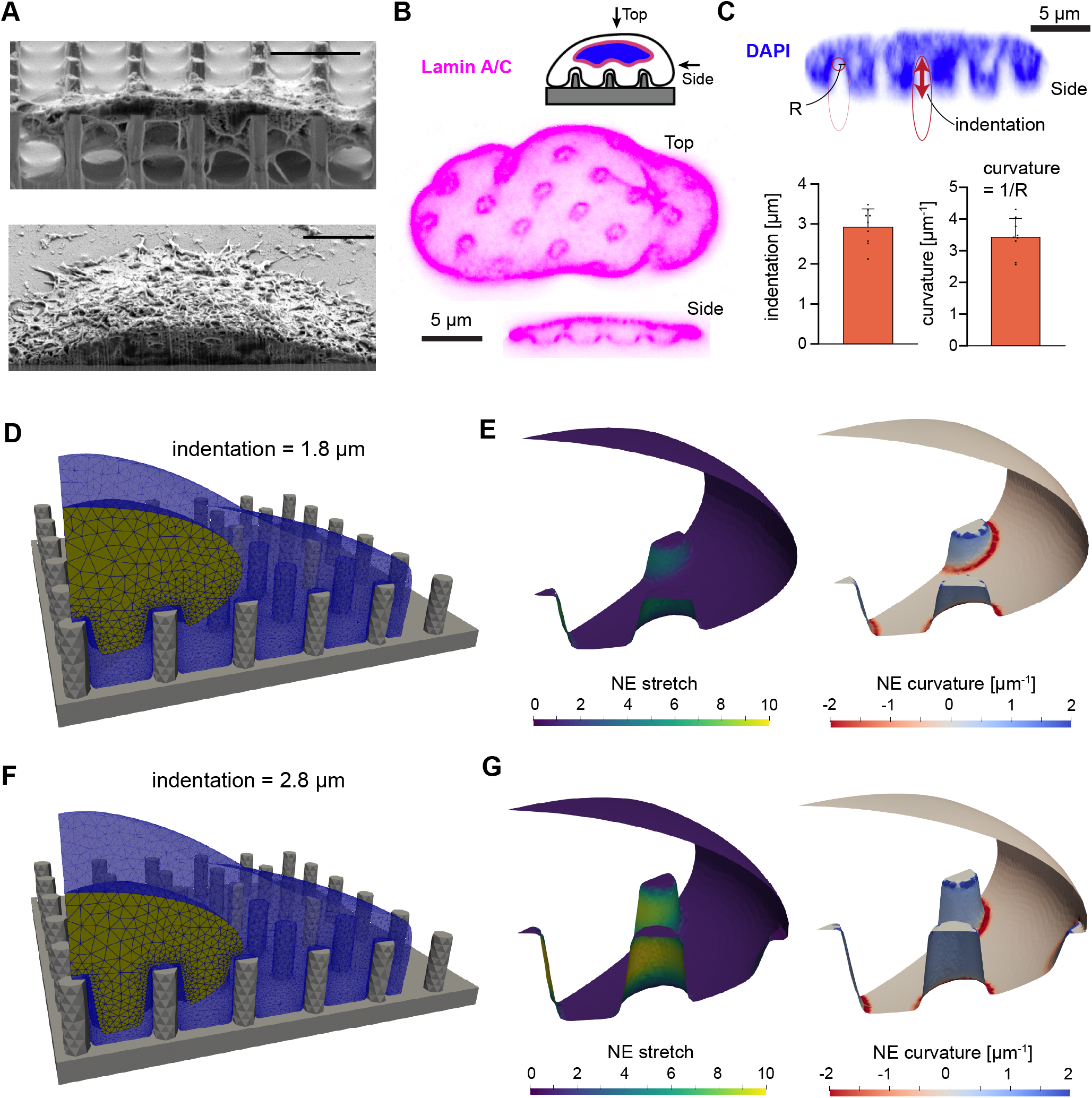
Plasma membrane nanotopography induces nanoscale curvature of the nuclear envelope. A) Crosssectional images of spread U2OS cells on a nanopillar substrate (upper) and flat surface (lower) using Focused Ion Beam Scanning Electron Microscopy. Scale bar denotes 5 µm. B) Confocal cross-section and reconstructed side view of lamin A/C within the NE. C) Quantification of nuclear indentation and NE curvature from reconstructed confocal microscopy images of DAPI. The circle of radius *R* and elongated ellipse represent the shapes drawn to quantify NE curvature and indentation, respectively. *N* = 9 in bar graphs, with each point representing the average indentation or curvature induced in a single cell nucleus. D-G) Meshes of model cells on substrates with *r*_*NP*_ = 500 nm, *h*_*NP*_ = 3 µm, *p*_*NP*_ = 3.5 µm and nuclear indentations of 1.8 µm (D) or 2.8 µm (G). Calculated NE stretch (Equation S47) and NE curvature (Equation S50) are displayed for each mesh in E and G.

We reproduced these nuclear deformations in our model by defining a deformation field on the surface of a spheroidal nucleus (Figure 3C-F). For simplicity, we assumed that material points were displaced mainly in the vertical (*z*) direction due to the nanopillar-induced stress. The resulting area stretch ratio, *α* (local ratio of deformed area to the initial area; i.e., area dilation), and curvature of the nuclear surface were then analytically calculated (Supporting Information, Section S5). We enforced conservation of nuclear volume by adjusting the lateral dimensions of the nucleus; i.e., the nucleus was wider upon indentation (Table S1). Regions of highest stretch were localized to the NE adjacent to the sides of the nanopillars, while the top surface remained mostly unstretched (Figure 3D,F). Importantly, these stretch ratios should not be taken to correspond to areal dilation of the nuclear membrane itself, but rather the elastic stretching of the underlying nuclear lamina.^[38]^ The spatially averaged mean curvature agrees well with experimental measurements; *H*_*PM*_ averaged over the membrane indented by the central nanopillar in Figure 3F is 3.4 µm^*−*1^.

### 2.4 Nuclear indentation enhances cytoskeletal activation and allows for increased YAP/TAZ transport through NPCs

We next measured the distribution of F-actin (Figure 4A) and Lamin A/C (Figure 4B) in U2OS cells on nanopillars. Cells were fixed 8 hours after seeding to allow sufficient time for adhesion and spreading on nanostructures. The focal plane was carefully chosen to ensure optimal visualization of the cellular structure and fluorescence images were taken that showed accumulation of F-actin around the nanopillars, in agreement with previous studies.^[17, 6]^ Lamin A/C also appeared to concentrate around nanopillars, which was at least in part attributed to a greater density of NE surface projected to the focal plane in cases of nuclear deformation.

**Figure 4.**
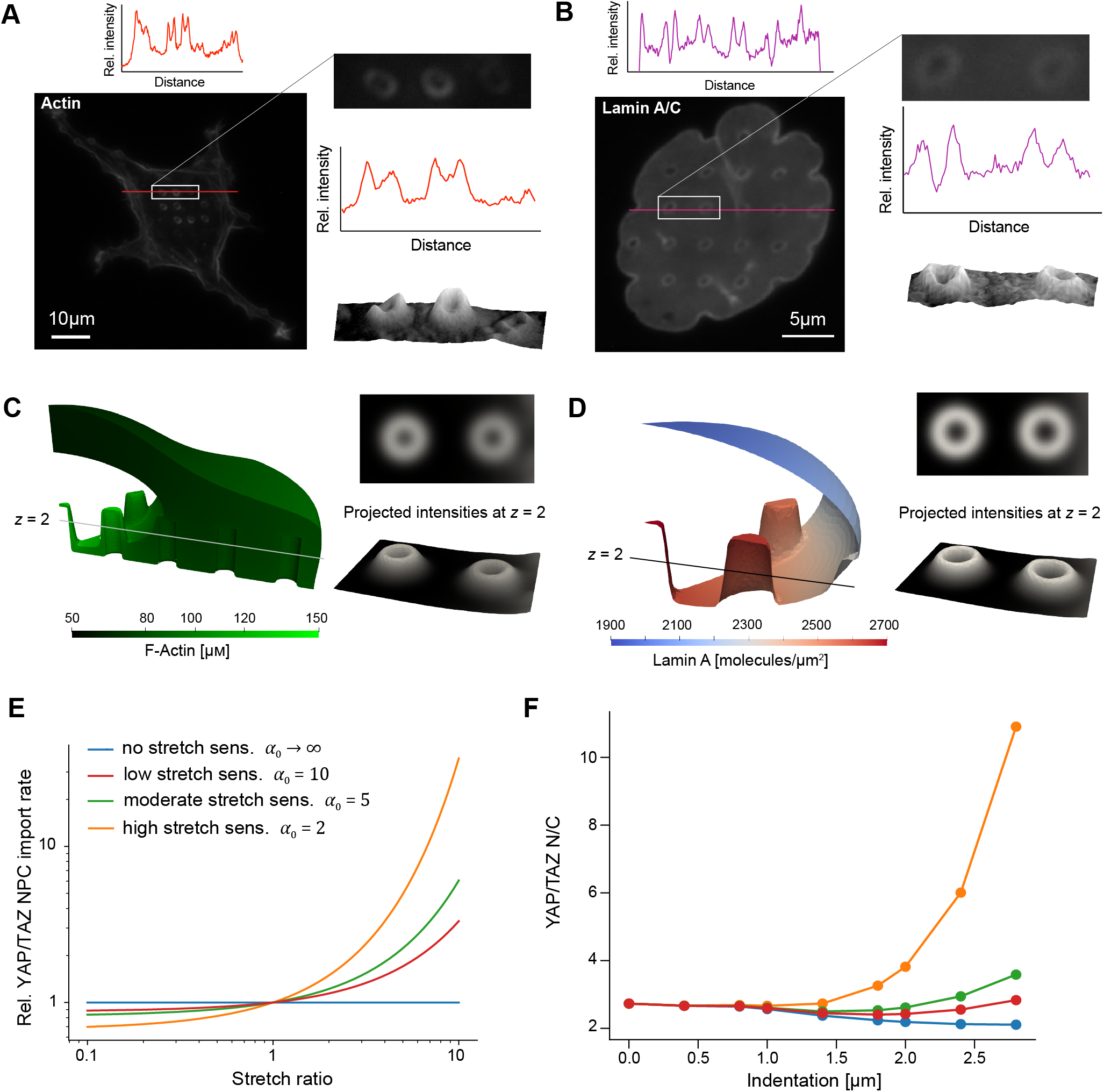
Effects of nuclear indentation on cytoskeletal activation, lamin organization, and nuclear transport. A) Fluorescence images of actin in a U2OS cell, with an intensity profile quantified along the red line. The right images show a magnified image of actin accumulation around 3 nanopillars, with the associated intensity profile (middle) and surface plot (lower). B) Fluorescence image of lamin A/C localization in a region of nuclear deformation on nanopillars, with the associated intensity profile along the red line. The right images show a magnified image of lamin intensity around 2 nanopillars, with the associated intensity profile (middle) and surface plot (lower). C) Actin distribution in a model cell at *t* = 10,000 s, with right images showing the projection to the *z* = 2 µm plane for comparison with fluorescence images. D) Distribution of non-phosphorylated lamin A in the NE at *t* = 10,000 s, with right images showing projected intensities at the *z* = 2 µm plane. E) Exponential relationship between YAP/TAZ import rate through NPCs (relative to the import rate without stretch sensitivity) and the stretch ratio of the NE for different values of *α*_0_. F) Steady-state YAP/TAZ N/C for different nuclear indentations.

We then simulated the effects of nuclear deformation on cytoskeletal activity and lamin phosphorylation in our computational model. Due to N-WASP-mediated actin polymerization near the surface of the nanopillars and the effects of confinement between the PM and NE, we observed a higher concentration of actin surrounding nanopillars in the region of nuclear deformation (Figure 4C). For comparison with experimental images, we computed the projected density of F-actin at *z* = 2 µm upon convolution with a Gaussian kernel for a cell at steady-state (see Methods, Equation (11)). This intensity map qualitatively matched experimental measurements (Figure 4A,C). Similarly, we examined the effects on lamin distribution, as indicated by the surface density of dephosphorylated lamin A. Phosphorylated lamin A was not included, as it is assumed to dissociate from the NE.^[34]^ In accordance with model assumptions, dephosphorylated lamin A was preferentially localized to regions with higher concentrations of F-actin, leading to higher lamin A densities near sites of nuclear indentation (Figure 4D). Once again, we computed the intensity projection onto the *z* = 2 µm plane, revealing a distribution quite similar to experimental measurements (Figure 3B,D). This local aggregation of lamin A agrees well with other measurements of lamin A distribution following cell deformation on nanoneedle substrates.^[39]^

Next, we predicted the effects of nuclear indentation on transport of YAP/TAZ through NPCs. Lamin A dephosphorylation directly potentiates NPC opening in our model, in cooperation with F-actin and activated myosin, under the assumption that stress fibers interact with the NE through the LINC complex to facilitate NPC stretching.^[33]^ In addition, previous measurements indicate that stretching of the NE (e.g., by nuclear compression or indentation) can directly facilitate YAP transport through NPCs.^[33, 35, 40]^ Therefore, we proposed that transport rate through NPCs exponentially scales with the local stretch ratio, *α*, of the NE (Figure 4E):

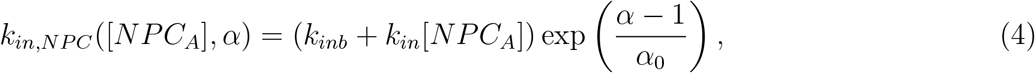

where *k*_*inb*_ and *k*_*in*_ are the basal and NPC-activation-dependent rates of YAP/TAZ transport into the nucleus, respectively, [*NPC*_*A*_] is the surface density of activated NPCs and *α*_0_ sets the sensitivity of NPC transport to stretch. According to this equation, entry is enhanced in regions of stretch (*α >* 1) and reduced in regions of surface compression (*α <* 1).

Examining the model across a range of nuclear indentations, we observed distinct elevations in YAP/TAZ N/C for increased indentation when *α*_0_ was less than or equal to 5 (Figure Figure 4F, Movies 7,8,9,10). In the absence of stretch sensitivity, the model surprisingly predicts reduced YAP/TAZ N/C at high levels of nuclear indentation. This occurs due to a reduced gap between the nucleus and PM at the top of the cell geometry, leading to increased cytoskeletal activation in the confined region between the NE and PM (Figure S2, Movie 7). In subsequent simulations, we fixed the stretch sensitivity at *α*_0_ = 5, as this value predicted an increase in YAP/TAZ N/C with nuclear indentation within the N/C range commonly measured experimentally.

Enhanced entry of YAP/TAZ at regions of high stretch, together with N-WASP-mediated actin assembly at closely apposed regions of PM, may compensate for weakened mechanosignaling due to endocytosis. This effect can be captured by considering a well-mixed limit of our model (derived in Section S3), in which YAP/TAZ N/C at steady-state is given by:

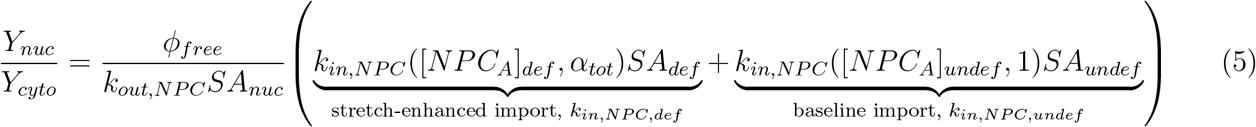

where *ϕ*_*free*_ is the fraction of free YAP throughout the cytosol, *α*_*tot*_ is the total stretch of the NE in the deformed region, *k*_*out,NPC*_ is the YAP/TAZ export rate through NPCs, *SA*_*def*_ and *SA*_*undef*_ are the surface area of the deformed and undeformed regions of NE (*SA*_*nuc*_ = *SA*_*def*_ + *SA*_*undef*_), and [*NPC*_*A*_]_*def*_ and [*NPC*]_*undef*_ are the average density of open NPCs in the deformed and undeformed regions. The effects predicted by this simplified model are plotted in Figure S3B, illustrating that NE stretch provides a tunable mechanism for changes in nuclear YAP/TAZ.

### 2.5 Nuclear rupture leads to high levels of YAP/TAZ in select cells on nanopillar substrates

In addition to the effects of NPC stretch due to nanopillar-induced deformation of the NE, recent experiments indicate that some cells seeded on nanopillar substrates experience transient NE rupture (NER).^[22]^ This was shown to cause a protein normally confined to the nucleus, Ku-80, to escape into the cytosol. We postulated that a similar effect might hold for YAP/TAZ; that is, NER might allow for increased nuclear entry and/or exit of YAP/TAZ (Figure 5A).

**Figure 5.**
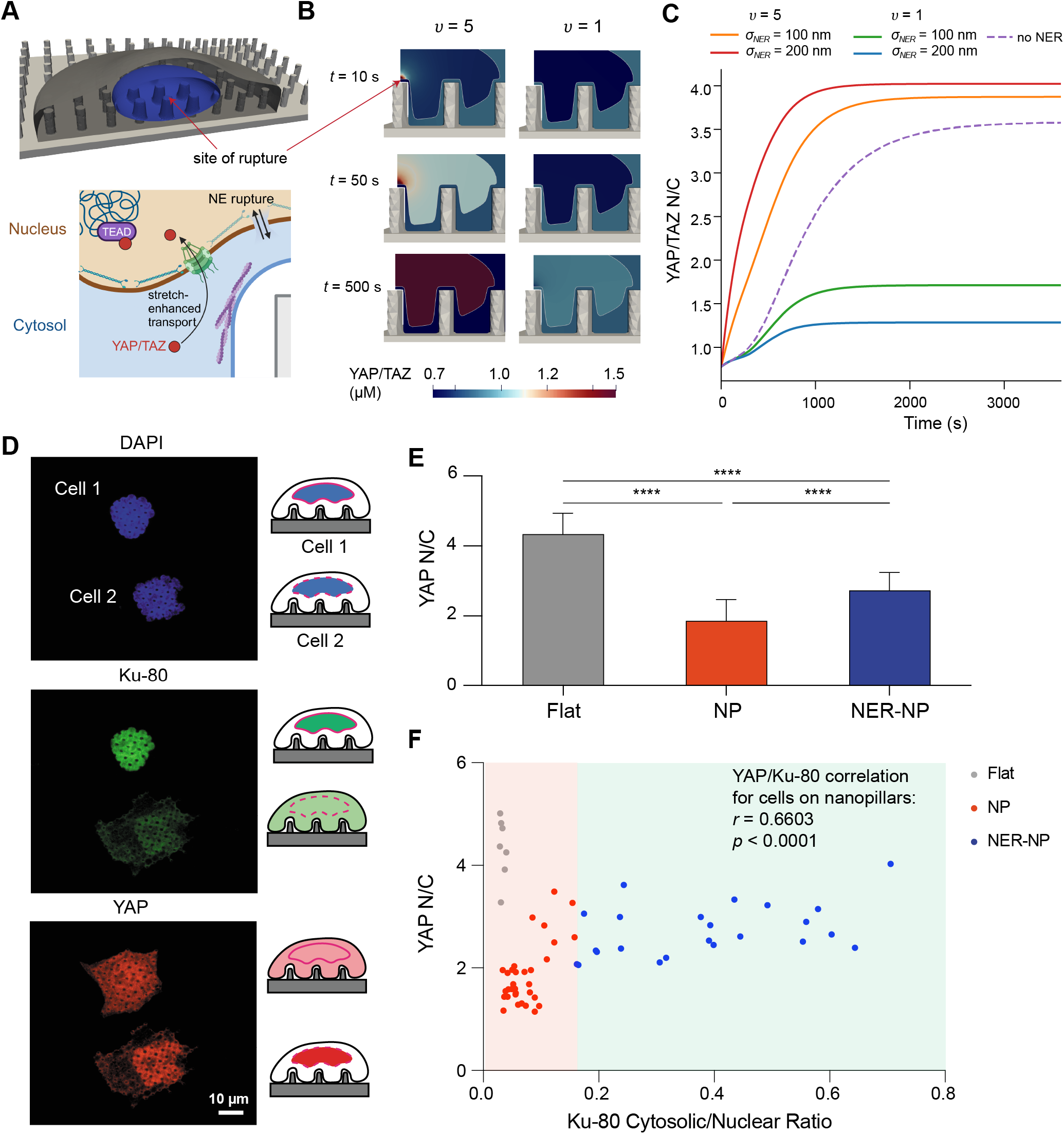
Model predictions and experimental measurements of YAP/TAZ transport following rupture of the nuclear envelope. A) Site of rupture in cell geometry (upper) and schematic depicting the combination of force-dependent NPC opening and molecular transport through NE rupture-induced pores (lower). B) Spatial distribution of YAP/TAZ in the cytosol and nucleus for *υ* = 5 and *υ* = 1. C) Dynamics of YAP/TAZ N/C for different pore radii (100 nm or 200 nm) and different values of *υ* (1 or 5), compared to simulations with no NER (dashed line), *k*_*rupture*_ = 100,000 µm^*−*2^s^*−*1^µm^*−*1^. D) Immunofluorescence imaging of cells stained for nuclear location (DAPI), NER marker (Ku-80), and YAP demonstrate mislocalization of Ku-80 to the cytosol and YAP to the nucleus in a cell exhibiting NER. E) Quantitative comparison of YAP N/C ratio of cells on flat or nanopillar substrates reveals higher translocation of YAP to the nucleus for cells exhibiting NER (NER-NP) compared to cells on nanopillars without NER (NP). \*\*\*\**p <* 0.001 according to two-way ANOVA followed by Tukey’s post-hoc test. Data is presented as mean ± standard deviation. F) Scatter plot illustrating the positive correlation between YAP N/C and Ku-80 cytosolic/nuclear ratio, where cells exhibiting NER show elevated YAP N/C. Correlation was assessed between YAP N/C and Ku-80 cytosolic/nuclear ratio, including all cells on nanopillar substrates. *r* denotes Pearson’s correlation coefficient and *p* is the *p*-value associated with the null hypothesis *r* = 0. In E and F, *N* = 7 for flat, *N* = 32 for NP, and *N* = 23 for NER-NP.

We therefore included an additional entry mechanism over a small patch of the nuclear surface. We consider a rupture-induced pore that opens rapidly at the central nanopillar, represented implicitly through a spatiotemporal function defining the relative permeability *p*_*NER*_ of the NE (Figure 5B):

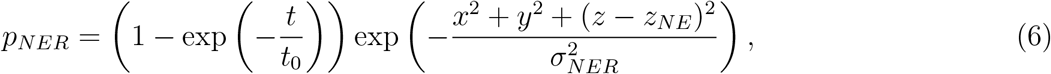

where *σ*_*NER*_ dictates the size of the pore and *z*_*NE*_ is the *z* value associated with the lower portion of the NE at *x* = 0, *y* = 0. Pore formation is initialized at *t* = 0, after which *p*_*NER*_ exponentially approaches 1 with a characteristic time of *t*_0_. We assume that the pore forms rapidly (*t*_0_ = 1 s), in line with previous measurements of membrane damage.^[41, 42]^

The net flux of YAP/TAZ is assumed to be diffusive, governed by the relation:

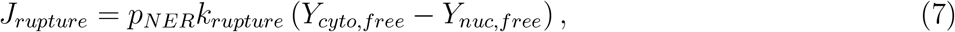

where *k*_*rupture*_ sets the magnitude of the transport rate (estimated assuming diffusive transport, Table S4) and *Y*_*nuc,free*_ is the free fraction of YAP/TAZ in the nucleus. Under the limit of rapid binding, this is proportional to the total YAP/TAZ in the nucleus:

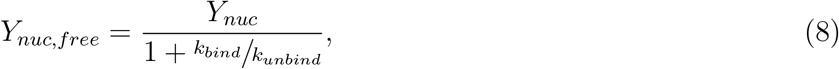

where *k*_*bind*_ and *k*_*unbind*_ are the rates of YAP/TAZ binding and unbinding to factors in the nucleus (e.g., TEADs), respectively. Writing *υ* as the ratio of total YAP/TAZ to free YAP/TAZ in the nucleus 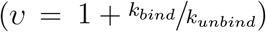 and substituting Equation (8) into Equation (7) yields:

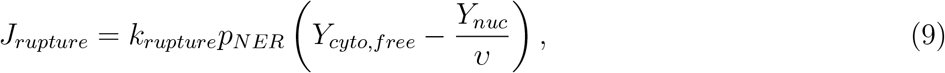

To test the effects of rupture-induced pore size and YAP nuclear retention in our model, we systematically varied *υ* and *σ*_*NER*_. Increasing *σ*_*NER*_ alters the contribution of transport through the rupture-induced pore relative to transport through NPCs, whereas *υ* determines the extent to which YAP/TAZ is retained in the nucleus. When *υ* = 5, NER caused increases in nuclear YAP/TAZ compared to conditions without rupture (Figure 5B-C, Movie 11). However, we found that when *υ* = 1, YAP/TAZ N/C is lower than the baseline condition (Figure 5B-C, Movie 12).

The overall contribution of NER on steady-state YAP/TAZ N/C can be assessed in a well-mixed framework, building on Equation (5):

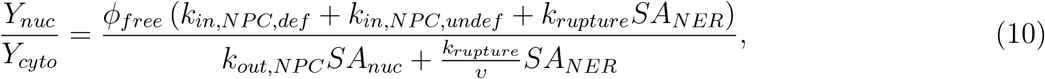

where *k*_*in,NPC,def*_ and *k*_*in,NPC,undef*_ are defined in Equation (5) and *SA*_*NER*_ is the surface are of the ruptureinduced pore 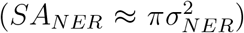. When transport through the rupture-induced pore dominates and the majority of YAP/TAZ is free for transport into the nucleus (*ϕ*_*free*_ → 1), YAP/TAZ N/C approaches *υ*. Furthermore, this expression predicts that nuclear YAP/TAZ N/C will increase due to NER only if *υ* is greater than YAP/TAZ N/C without rupture, as plotted in Figure S3C.

To test these model predictions, we examined the YAP N/C for cells on nanopillars in our experiments. As recently reported,^[22]^ some cells showed evidence of NER, indicated by the cytosolic mislocalization of Ku-80 (Figure 5D). We observed an increase in nuclear localization of YAP in the same cells (Figure 5D). We then examined the average behavior over many cells, defining cases of NER as those cells with Ku-80 cytosolic-to-nuclear ratios greater than 0.16, as in our recent study^[22]^ In agreement with previous studies,^[14, 20]^ our quantitative analysis confirmed that, among cells not experiencing NER, those on flat substrates exhibited significantly higher YAP N/C ratio compared to those on nanopillars (Figure 5E). Additionally, cells exhibiting NER had a significantly elevated YAP N/C ratio compared to cells on nanopillars without rupture (Figure 5E), suggesting NER enhances the translocation of YAP to the nucleus. This is further supported by the statistically significant positive correlation between the cytosolic-to-nuclear Ku-80 ratio and YAP N/C for cells on nanopillar substrates (Figure 5F).

In relation to our model, this data indicates that *υ* must be greater than the highest YAP N/C in ruptured cells (≈4), implying that less than 25% of YAP in the nucleus is free to diffuse (Equation (8)). This agrees well with the slow effective diffusion of YAP measured in the nuclear volume.^[43]^ Together, our experiments and simulations position NER as an additional mechanism for YAP/TAZ transport across the NE in response to substrate topography.

## Discussion

Mechanotransduction, the process of cell signaling in response to mechanical cues, is a key regulator of cell state, proliferation, and differentiation.^[44, 2]^ One key mechanical cue is the local stiffness of the pericellular microenvironment, which regulates how cells sense their immediate environment.^[24, 26, 45]^ Our findings highlight the additional importance of nanotopography in cell mechanosignaling. Many recent experiments demonstrate the importance of nano-scale topographical cues to local cytoskeletal assembly,^[16, 5, 6]^ clathrin-mediated endocytosis,^[14, 13]^ and deformation of the NE,^[21, 22, 20]^ each of which have significant downstream effects on the nuclear translocation of YAP/TAZ. However, our computational model is the first to incorporate these effects into a predictive computational model of cell mechanosignaling. Our findings reveal that cell signaling integrates many mechanical and geometric cues including PM curvature, cytoskeletal assembly, and nuclear mechanics, resulting in mechanoadaptation.

Many previous models of YAP/TAZ mechanotransduction assume the cell to be a composite of well-mixed volumes.^[23, 25, 46]^ This results in systems of ordinary differential equations (ODEs), which are not generally equipped to handle the complex dynamics of signaling networks in realistic cell geometries. Other recent models consider the spatial dynamics of mechanotransduction,^[24, 27, 47, 48]^ but these simulations were restricted to relatively simple model geometries. Here, we consider detailed cell geometries on several different nanopillar substrates, making use of our new software package, SMART,^[28, 27]^ which is especially well-suited for simulating realistic cell geometries.

Simple extrapolation from past modeling work^[24, 27]^ might lead one to expect that cells on substrates with nanoscale curvature exhibit enhanced cytoskeletal and YAP/TAZ activation due to increased PM surface area to cytosolic volume ratios near the substrate. However, the effect of nanotopography on mechanotransduction signaling in experiments contradicts this prediction; while nanotopography does induce local actin assembly^[17, 16]^ and may potentiate YAP/TAZ transport through NPC stretching,^[33, 40]^ the dominant effect appears to be attenuation of cytoskeletal assembly on nanopillar substrates due to endocytosis of β_1_ integrins.^[14]^

Here, we integrate these confounding effects in a general model of cell response and adaptation to nanotopography (Figure 6). Alongside purely geometric effects, we posit that cytoskeletal activity is globally inhibited due to integrin endocytosis,^[14]^ but locally enhanced via curvature-dependent activation of N-WASP.^[17, 16]^ Global inhibition leads to lower levels of nuclear YAP/TAZ as a function of nanotopographical features such as nanopillar density (Figure 6A). We also demonstrate a crucial role for YAP/TAZ transport as a function of NE stretch and rupture. Increased nuclear indentation by nanopillars is predicted to increase the rate of YAP/TAZ import in a stretch-dependent manner (Figure 6B). NER was also shown to influence YAP/TAZ N/C (Figure 6C); when YAP/TAZ is retained within the nucleus due to binding to nuclear factors, our model predicts YAP/TAZ nuclear accumulation, validated by experimental measurements. The overall balance between local activation and global inhibition (Figure 6D) is reminiscent of proposed mechanisms for cell navigation in response to chemical gradients,^[49, 50]^ suggesting that mechanoadaptation described by our model may be well-suited to steer migrating cells through complex topographies.

**Figure 6.**
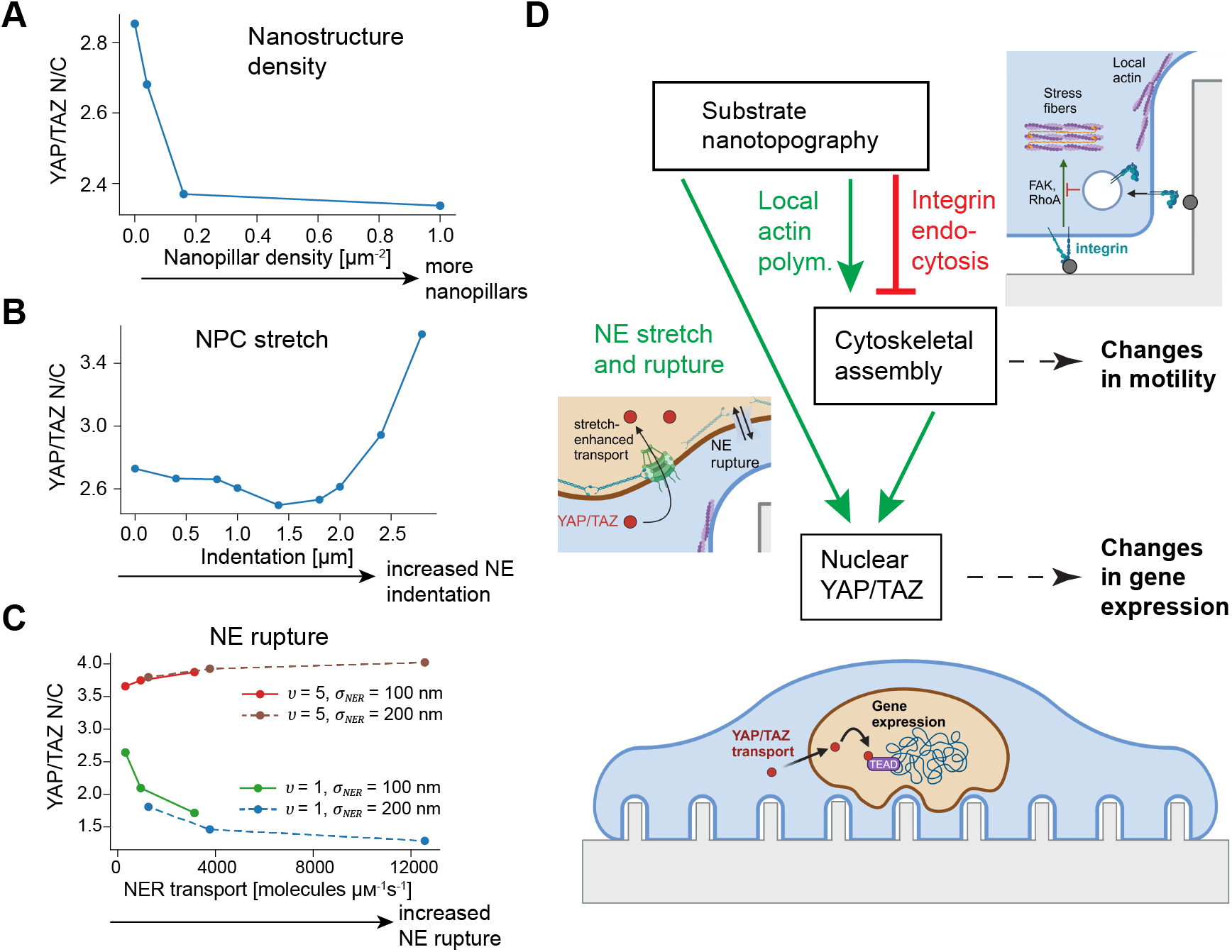
Mechanisms for mechanoadaptation to nanotopography. A-C) Summary of the effects of nanopillar spacing/density (A; plotted for *r*_*NP*_ = 100 nm), NPC stretch due to NE indentation with *α*_0_ = 5.0 (B), and NER (C) on YAP/TAZ N/C at *t* = 3600 s. In panel C, NER transport rate is defined as 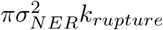, with *k*_*rupture*_ tested at 10,000, 30,000, and 100,000 s^*−*1^ µm^*−*1^ µm^*−*2^ for each condition. D) Schematic summarizing the effects of nanotopography on YAP/TAZ mechanotransduction in our model, including negative regulation by integrin endocytosis in parallel with positive cues due to local actin polymerization, NE stretch, and NER.

YAP/TAZ transport through rupture-induced pores is assumed to be governed by passive diffusion in our model, in contrast to active transport through NPCs. The size of such ruptures and their formation and repair dynamics remain unclear. The fact that Ku-80 cytosolic-to-nuclear ratios span a wide range of different values (Figure 5F) implies that cells exhibit different extents of nuclear rupture, likely due to nuclear repair over the timescale of about 1 hour.^[22]^ The process of NE rupture and repair require additional consideration in future experiments and models and may well influence the transport of other molecules in addition to YAP/TAZ. Such work can be facilitated by novel experimental tools such as FRET-based lamin A/C tension sensors.^[51, 52]^

While we just recently reported NER events in response to nanotopographical features,^[22]^ related transient changes in NE integrity have been observed in response to different mechanical stimuli during cancer and immune cell migration.^[53, 54]^ NE disruption is generally understood to be detrimental to cell function, associated with diseases such as laminopathies, cardiomyopathies, and cancer, or with cellular aging (senescence).^[55, 56]^ However, our findings suggest that at least some cases of temporary NE rupture can enhance cell mechanotransduction. Whether such responses are observed in vivo under non-pathological conditions remains to be seen.

N-WASP-mediated actin assembly around nano-curved structures, stretch-activated NPC opening, and rupture of the NE are all potential compensatory mechanisms for reduced signaling due to integrin endocytosis at sites of inward curvature.^[13, 14]^ (Figure 6D) The contribution of each process to YAP/TAZ signaling depends on adjacency between the PM and NE (see Figure S2) and on the extent of nuclear deformation. Experiments and models indicate that the nucleus is deformed due to combined effects of downward force of the perinuclear actin cap pushing downward on the nucleus and to the anchoring of the lower NE to the PM.^[21, 57]^ This process of nuclear deformation therefore effectively acts as an internal dial by which the cell can tune nuclear transport of YAP/TAZ. This may be especially important for cancer cells, such as the U2OS cells tested here, as changes in nuclear YAP/TAZ are a key determinant of differentiation and metastasis across many cancers.^[2, 58]^

The curvature- and deformation-dependent signaling events explored in this paper apply well beyond YAP/TAZ mechanotransduction networks. First, the modulation of cytoskeletal assembly by PM curvature has implications for many cell responses to different nanotopographies, from immune cell migration^[5]^ to alignment of fibroblasts along collagen fibrils.^[59]^ Furthermore, the enhancement of endocytosis at sites of high curvature could have effects on many different signaling cascades initiated at the PM. Similarly, NPC stretching and NER could influence the transport of many different factors important to mechanotransduction to and from the nucleus in addition to YAP/TAZ, such as myocardin-related transcription factors^[60]^ or Twist1.^[61]^ Thus, the integration of mechanical and chemical cues at multiple length scales is critical for mechanoadaptation.

## 3 Methods

### 3.1 Mesh generation

All meshes were generated using the open-source mesh generator, Gmsh.^[36]^ For simplicity, it is assumed that nanopillar substrates are well-described by a composite of a flat surface and cylindrical structures of a specified height, spacing (pitch), and radius. Furthermore, model cell geometries formed a circular contact area over the substrate, with one nanopillar positioned directly at the center of the contact region. In this case, the model geometry cell has four unique symmetry axes (*x* = 0, *y* = 0, *y* = *x*, and *y* = *−x*), allowing a reduction in computational cost by simulating 1/8 of the full geometry. The PM closely conformed to the nanopillar surface, in agreement with experimental observations.^[21]^ In particular, the PM was fixed 50 nm from the nanopillar surface in all locations except close to the lower flat substrate, where the membrane curved to smoothly meet the surface with a characteristic radius of 200 nm (Figure S4A, Section S4). The projected cell-substrate contact area was prescribed according to experimental measurements for each condition, with the added constraint of conserved cell and nuclear volume matching previous measurements of U2OS cells.^[62]^

The nuclear mesh was continuous with the cytosolic mesh, with its reference boundary specified as an oblate spheroid with aspect ratio 2.21:1 (Equation S5) and volume matching that measured in experiments with U2OS cells.^[62]^ The bottom of the nucleus was positioned 0.2 µm above the PM at the central nanopillar. In cases where the top of the nucleus did not fit within 0.8 µm of the top PM boundary, the vertical axis length was reduced and the radial axis was expanded to conserve volume (Equations S8 and S7). In cases of nuclear indentation, the nuclear boundary was modified in Gmsh according to Equation S38 as described in Section S5.

A summary of geometric parameters, volume and area of each subsurface and subvolume, and number of nodes and elements in each subsurface and subvolume are given for each mesh in Table S1. Details of mesh generation are further discussed in Section S1. Meshes used in the final simulations are freely available via Zenodo.^[63]^

### 3.2 Spatial simulations of YAP/TAZ signaling

All simulations were conducted using the Python package, Spatial Modeling Algorithms for Reaction and Transport (SMART).^[28, 27]^ The SMART framework takes in user specifications of the species, reactions, parameters, and compartments involved in a given reaction-transport network. This model involves 15 species across four compartments: the PM, the cytosol, the NE, and the nucleoplasm (Table S2). All reactions between species and parameters governing these reactions are summarized in Tables S3 and S4. Other than the equations given in the main text, all reactions were used as specified in previous work.^[24, 27]^

These specifications, together with diffusion coefficients and initial conditions for each species, define the system of nonlinear mixed dimensional PDEs governing the system (Equations S10-S24). SMART internally converts these equations to their variational form and assembles a monolithic finite element system using the finite element software package, FEniCS. Linear finite element shape functions are used for each triangular or tetrahedral element and the system is discretized in time according to a backward (implicit) Euler scheme. The nonlinear system is solved at each time step using Newton-Raphson iterations in PetSc.^[64]^ The initial time step was 0.01 s, after which SMART adaptively adjusted the time step according to the number of Newton iterations required for convergence at the previous time. Code for model specifications, mesh creation, and data analysis is freely available on Github.^[65]^ Results files are also available via Zenodo.^[63]^ All data was visualized in Paraview.

### 3.3 Computation of projected intensity maps from simulation data

For comparison with experimental images, we computed a projection of our finite element solution onto the *z* = 2 plane in Figure 4. To compute this, the spatial solution was convolved with a Gaussian kernel; that is, the projected intensity *I* was given by:

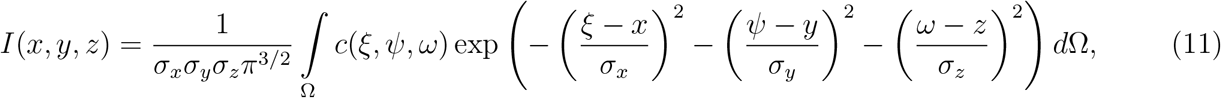

where *c*(*x, y, z*) is the spatially defined finite element solution for lamin A or F-actin, and *σ*_*x*_ = *σ*_*y*_ = 0.2 µm and *σ*_*z*_ = 0.4 µm to represent the effects of blurring in the *x*-*y* plane vs. blurring in the *z* direction. This mimics the effect of convolution with the point-spread function (PSF) to compute an image in optics. Here, a simpler functional form than a conventional PSF is used as the comparison with fluorescence images is purely qualitative. Note that the domain Ω corresponds to either the cytosol or the NE in the case of F-actin and lamin A, respectively.

### 3.4 Nanopillar substrate fabrication and preparation

Nanopillars were fabricated as previously described.^[12]^ Briefly, quartz wafers were cleaned and spin coated with AZ 1512 photoresist (EMD Performance Materials) with 1.2 µm thickness. Patterning and exposure were performed using the Heidelberg maskless aligner (MLA) with a 375 nm laser at 300 mJ/cm^2^. Chromium (Cr, 99.998%) was deposited using the Temescal Ebeam Evaporator, followed by lift-off. Reactive ion etching (RIE) with Ar and CHF_3_ gases was performed over a 50-minute process using the Oxford Plasmalab 80plus. Finally, wet etching with Cr etchant (Transene Company Inc) and buffered oxide etch (KMG Chemicals Inc., USA) 20:1 completed the fabrication, and the wafers were diced into 1 cm^2^ chips for further experimentation. Chips were sterilized with 70% ethanol, rinsed with deionized water, and air-dried. They were then UVO-treated for 10 minutes and incubated with 100 µg mL^*−*1^ poly-L-lysine (Sigma-Aldrich, USA) solution for 30 minutes. After washing with 1X phosphate buffered saline (PBS; Gibco, USA), the chips were treated with 0.5% glutaraldehyde for crosslinking, followed by another PBS wash. Finally, a pre-warmed 0.1% gelatin solution (Sigma-Aldrich, USA) was applied, and the chips were incubated at room temperature for 30 minutes.

### 3.5 U2OS cell experiments

U2OS (ATCC) were cultured in McCoy’s 5A medium (ATCC) supplemented with 10% fetal bovine serum (FBS; Invitrogen, USA) and 1% penicillin-streptomycin (PenStrep; Thermo Fisher Scientific, USA) and were incubated at 37°C with 5% CO_2_. Cells were seeded onto nanopillar chips by first detaching them with TrypLE Express Enzyme (Gibco, USA) and preparing a 100 µL suspension containing 50,000 cells. The cell suspension was then dropped onto the chips and allowed to adhere for 10 minutes. After cell adhesion, 1 mL of growth medium was added to the 24-well plate, and the cells were incubated at 37°C and 5% CO_2_ for further analysis.

### 3.6 Immunostaining

Cells were fixed after 8 hours of incubation using 4% paraformaldehyde (Electron Microscopy Sciences, USA) at 37°C for 10 minutes. Then cells were permeabilized with Triton X-100 (Sigma-Aldrich, USA) for 10 minutes and blocked in 1% Bovine Serum Albumin (BSA) (Thermo Fisher Scientific, USA) overnight before immunostaining. Rabbit anti-Ku-80 (Cell Signaling Technology, USA) at 1:200 dilution and mouse Anti-YAP1 Antibody (63.7) (Santa Cruz Biotechnology, USA) at 1:100 dilution in PBS were used for primary antibody staining overnight at 4°C. Samples were washed 10 times with PBS to remove all primary antibodies. Samples were then incubated with secondary antibodies conjugated with Alexa Fluor 488 (anti-rabbit IgG; Thermo Fisher Scientific, USA) or Alexa Fluor 594 (anti-mouse IgG; Thermo Fisher Scientific, USA) at 1:1000 dilution for 1 hour at room temperature. Samples were washed in PBS and stained with 4*′*,6-diamidino-2-phenylindole (DAPI) (Thermo Fisher Scientific, USA) for 5 minutes. For the actin- and lamin-labeled cells shown in Figure 4, the same protocol was utilized, instead using a single primary antibody (mouse anti Lamin A/C from Biolegend at 1:400 dilution) and a single secondary antibody (anti-mouse IgG conjugated with Alexa Fluor 647 at 1:1000 dilution; Thermo Fisher Scientific, USA), followed by staining with Alexa 594-phalloidin (InvitrogenTM, USA) for 20 minutes.

### 3.7 FIB-SEM

Preparation of samples for FIB-SEM imaging were adapted from methods previously described.^[66]^ The live cells were first fixed in 2.5% glutaraldehyde (Sigma-Aldrich, USA) overnight at 4°C, followed by 3 rinses in PBS pH 7.4 (Gibco, USA). Samples were then rinsed with autoclaved DI water and dehydrated using a series of increasing ethanol (Thermo Scientific, USA) dilutions of 30, 50, 70, 80, 90, and 100 percent, and incubated for a period of 10 minutes per step. The ethanol was then displaced from each sample by covering the chip with a 1:1 v/v of pure ethanol to Hexamethyldisilazane (HMDS) (Sigma-Aldrich, USA) for 15 minutes, followed by incubation in 100% HDMS for an additional 15 minutes. The HMDS was then removed from the sample and the sample was air-dried overnight in a chemical fume hood. The dried samples were then sputter-coated with an iridium oxide layer, mounted on SEM stubs, and introduced into an FEI Scios DualBeam system (FIB/SEM). Cross-sectioning was performed using a 5 nA ion beam current, and a 0.5 nA beam current was used for cleaning the cross section prior to imaging. A 45° tilt was applied to the microscope stage to enhance the observation of nanopillar-NE interaction and high-resolution images were acquired at 5 kV using an Everhart-Thornley Detector (ETD).

### 3.8 Fluorescence microscopy

Widefield fluorescence images were captured using an Echo Revolve microscope equipped with a 40× PLAN Fluorite LWD CC objective with numerical aperture 0.60. Confocal imaging was performed on a Nikon AXR microscope with a four-line laser system (405, 488, 561, and 640 nm) mounted on a Nikon Ti2 platform. Image stacks were captured in galvano mode with unidirectional scanning. *z*-stacks were acquired with 0.175 µm steps to capture the whole cell.

### 3.9 Image analysis

We utilized ImageJ version 1.53 (National Institutes of Health, USA) for image analysis. To quantify curvature, confocal images were reconstructed from side views using the reslice feature of ImageJ software. The largest fitting ellipse was manually drawn in an indented region of the nucleus, and the radius of a fitting circle tangent to the upper portion of this ellipse was calculated using ImageJ software (Figure 3C). The nuclear indentation was defined as the distance from the top of the ellipse to its center (Figure 3C).

To assess NE rupture and YAP localization, immunofluorescence staining was used to distinguish between nuclear and cytoplasmic regions, with the nucleus defined using a threshold based on the DAPI channel. Ku-80 and YAP intensity was measured within these nuclear boundaries, and cytoplasmic intensity was quantified in areas outside the nucleus but within the cell borders. Line profile analysis was performed using the “Plot Profile” function and surface profile analysis using the “Surface Plot” function in ImageJ.

### 3.10 Statistical analysis

For statistical analysis, we used GraphPad Prism (v9, GraphPad Software). Data for YAP nuclear to cytoplasmic ratio were analyzed using two-way ANOVA followed by Tukey’s post-hoc test. * denotes p *≤* 0.05, ** denotes p *≤* 0.01, and *** denotes p *≤* 0.001.

## Supporting Information

Supporting Information is available from the Wiley Online Library or from the author.

## Acknowledgements

The authors would like to thank Dr. Marie Rognes at Simula Research Laboratory for her support and advice throughout the development of SMART that led to these simulations. We are also grateful for helpful feedback on our manuscript from Dr. Nick Bergkamp and Dr. Aravind Chandrasekaran, Kyle Stark, and Ji Yeon Hyun at UCSD. Support in code development for SMART and guidance for mesh generation via Gmsh was provided by Dr. Jørgen Dokken and Dr. Henrik Finsberg at Simula Research Laboratory.

Simulation results presented in this paper benefited from the Triton Shared Compute Cluster at the San Diego Supercomputer Center.^[67]^ This work was performed in part at the San Diego Nanotechnology Infrastructure (SDNI) of UCSD, a member of the National Nanotechnology Coordinated Infrastructure, which was supported by the National Science Foundation (Grant ECCS-2025752). The authors acknowledge help from Dr. Zhicheng Long for conducting FIB-SEM. This work was supported in part by the Wu Tsai Human Performance Alliance at the University of California, San Diego (to P.R.) and by NIH R01GM132106 (to P.R.). E.A.F. is supported by the National Science Foundation under grant EEC-2127509 to the American Society for Engineering Education and by the Wu Tsai Human Performance Alliance. This work was additionally supported by the Air Force Office of Scientific Research YIP award (Award Number: 311616-00001), the Cancer Research Coordinating Committee faculty seed grant, National Science Foundation (Grant DMR-2011924) to Z.J., and National Science Foundation (Grant DMR-2011924) to Z.J. and A.S. P.R. is a consultant for Simula Research Laboratories in Oslo, Norway and receives income. The terms of this arrangement have been reviewed and approved by the University of California, San Diego in accordance with its conflict-of-interest policies.

## Data availability statement

The code that supports the findings of this study is openly available in Zenodo athttps://doi.org/10.5281/zenodo.13952739.^[65]^ The data that support the findings of this study are openly available in Zenodo at https://doi.org/10.5281/zenodo.13948827.^[63]^

## Author contributions

E.A.F.: Conceptualization, Data Curation, Formal Analysis, Investigation, Methodology, Software, Visualization, Writing Original Draft, Review and Editing; E.S.: Conceptualization, Data Curation, Formal Analysis, Investigation, Methodology, Writing Original Draft, Review and Editing; V.P.: Investigation, Methodology; D.P.M.: Investigation, Methodology; L.S.: Investigation, Methodology, Review and Editing; Z.J.: Conceptualization, Funding Acquisition, Resources, Project Administration, Supervision, Review and Editing; P.R.: Conceptualization, Funding Acquisition, Resources, Project Administration, Supervision, Review and Editing

