## Supplementary Text, Figures, and Tables for "Nanoscale Curvature Regulates YAP/TAZ Nuclear Localization Through Nuclear Deformation and Rupture"

### Nanoscale curvature of the plasma membrane regulates mechanoadaptation through nuclear deformation and rupture, Supplementary Information

Emmet A. Francis[1,2,*], Einollah Sarikhani[3,*], Vrund Patel[3], Dhivya Pushpa Meganathan[3], Leah Sadr[3], Zeinab Jahed[3,**], and Padmini Rangamani[1,2,**]

[1]Department of Pharmacology, University of California San Diego, La Jolla, CA, USA
[2]Department of Mechanical and Aerospace Engineering, University of California San Diego, La Jolla, CA, USA
[3]Department of Nanoengineering, University of California San Diego, La Jolla, CA, USA
[*]these two authors contributed equally to this work

#### S1 Mesh generation for cells on nanopillar substrates

Cell geometries on substrates with nanopillars were generated in Gmsh [1]. As a reference geometry, we used a vertically shifted version of the axisymmetric cell shape previously defined [2], for which the plasma membrane surface satisfies:

$$\begin{aligned} \left(1 - \frac{(z-h_{NP})^4}{a+(z-h_{NP})^4}\right)(r^2+(z-h_{NP})^2) + \frac{b(r^2+((z-h_{NP})+c)^2)(z-h_{NP})^4}{d+(z-h_{NP})^4} &= R_c^2 \qquad z \geq h_{NP}, \\ r &= R_c \qquad 0 \leq z < h_{NP}, \end{aligned} \tag{S1}$$

where $r$ and $z$ are cylindrical polar coordinates, $R_c$ is the cell-substrate contact radius, and $a, b, c,$ and $d$ are constants adjusted to conserve cell volume for a given contact area and the spacing, height, and radius of nanopillars. The cytosolic volume was computed as:

$$V_{cyto} = V_{shape}(a,b,c,d,R_c) + \pi R_c^2 h_{NP} - V_{NP}(r_{NP}, p_{NP}, h_{NP}, R_c) - V_{nuc}, \tag{S2}$$

where $V_{shape}(a,b,c,d,R_c)$ is the volume of the shape above the nanopillars. We estimated this volume by computing the volume for a discretized version of the contour; given the list of $r, z$ coordinates over the contour ($\mathbf{r} = [r_1, r_2, ..., r_N], \mathbf{z} = [z_1, z_2, ..., z_N]$), the volume is:

$$\begin{aligned} V_{shape} &= \sum_{n=1}^{N-1} \pi \int_{z_n}^{z_{n+1}} \left(r_n + \frac{r_{n+1}-r_n}{z_{n+1}-z_n}(z-z_n)\right)^2 dz \\ &= \sum_{n=1}^{N-1} \pi(z_{n+1}-z_n)\left(r_n^2 + r_n(r_{n+1}-r_n) + \frac{1}{3}(r_{n+1}-r_n)^2\right), \end{aligned} \tag{S3}$$

$V_{NP}(r_{NP}, p_{NP}, h_{NP}, R_c)$ is the volume excluded by the nanopillars over the contact region (see Figure S4B for annotated volume regions):

$$V_{NP}(r_{NP}, p_{NP}, h_{NP}, R_c) = N_{NP}\pi \Bigg[ \underbrace{d_{steric}\left( r_{NP}^2 + r_{NP} d_{steric} \frac{\pi}{2} + 2\frac{d_{steric}^2}{3} \right)}_{\text{volume 1 (top)}} + \underbrace{(r_{NP} + d_{steric})^2 (h_{NP} - d_{steric} - d_{curv})}_{\text{volume 2 (middle)}} + \underbrace{d_{curv}\left( (r_{NP} + d_{steric} + d_{curv})^2 - (r_{NP} + d_{steric} + d_{curv}) d_{curv} \frac{\pi}{2} + 2\frac{d_{curv}^2}{3} \right)}_{\text{volume 3 (bottom)}} \Bigg], \tag{S4}$$

where $N_{NP}$ is the number of nanopillars with which the cell intersects according to its specified contact area. For simplicity, we do not consider any nanopillars that would intersect the cell at the boundary of the contact region; that is, $N_{NP}$ must be a whole number.

To determine the boundary coordinates defined by implicit boundaries in Equation (S1), we solved for $r$ at each $z$ value using `solveset` in Sympy [3]. For a given contact area $A_c$, constants $a, b, c, d$ were adjusted iteratively until the volume was within 1% of the target volume. Small normal displacements of boundary nodes were then applied to adjust the boundary to within 0.1% of the appropriate volume. The U2OS total cell volume (including the nucleus) was fixed at 4,000 $\mu m^3$ assuming the resting cell radius is about 10 μm [4].

In the absence of indentation, the nuclear envelope (NE) forms an oblate spheroid:

$$\left(\frac{r}{a_{nuc}}\right)^2 + \left(\frac{z - z_{nuc}}{b_{nuc}}\right)^2 = 1, \tag{S5}$$

such that the nuclear volume is:

$$V_{nuc} = \frac{4}{3}\pi a_{nuc}^2 b_{nuc}. \tag{S6}$$

The center of the nucleus was placed at $z_{nuc} = h_{NP} - \Delta z_{indent} + 0.2\ \mu m + b_{nuc}$, where $\Delta z_{indent}$ is the nuclear deformation in the z-direction at the central nanopillar. In cases where the top of the nucleus was within 0.8 μm of the upper plasma membrane, $a_{nuc}$ and $b_{nuc}$ were adjusted while conserving nuclear volume:

$$b_{nuc,new} = \frac{z_{max} - h_{NP} + \Delta z_{indent} - 1.0}{2}, \tag{S7}$$

$$a_{nuc,new} = \sqrt{\frac{3V_{nuc}}{4\pi b_{nuc,new}}}, \tag{S8}$$

where $z_{max}$ is the $z$ value of the upper plasma membrane at $r = 0$. We constrained the nuclear volume by considering data from Koch et al. [5], in which U2OS cells were confined in microtubes and the dimensions of their nuclei were measured. Approximating the nuclei as ellipsoids, the average volume measured across microtube diameters was 550 $\mu m^3$; therefore, we fixed nuclear volume at this value in our simulations. In the cases shown in Fig S2, the nucleus is shifted vertically by setting $\Delta z_{indent}$ to a negative value. Equations governing the shape of the nucleus indented by nanopillars (that is, positive values of $\Delta z_{indent}$) are given in Section S5 below.

All resulting mesh statistics are summarized in Table S1.

**Table S1:** Mesh statistics.

| | Constants (in µm) | Vertices | Cells | Size |
|---|---|---|---|---|
| | $r_{NP} = 0.1, h_{NP} = 1.0, p_{NP} = 5.0, R_c = 20.25$ | | | |
| Cyto | $a = 2885, b = 0.4428, c = 24.37, d = 10.47$ | 14965 | 67920 | $4.367 \times 10^2$ µm$^3$ |
| PM | | 4596 | 8789 | $3.560 \times 10^2$ µm$^2$ |
| Nuc | $a_{nuc} = 7.740, b_{nuc} = 2.198, z_{nuc} = 3.398$ | 2652 | 8928 | $6.858 \times 10^1$ µm$^3$ |
| NE | | 1923 | 124 | $5.450 \times 10^1$ µm$^2$ |
| | $r_{NP} = 0.1, h_{NP} = 1.0, p_{NP} = 2.5, R_c = 18.52$ | | | |
| Cyto | $a = 2588, b = 0.4311, c = 20.18, d = 15.84$ | 23987 | 114150 | $4.364 \times 10^2$ µm$^3$ |
| PM | | 7472 | 14341 | $3.201 \times 10^2$ µm$^2$ |
| Nuc | $a_{nuc} = 6.625,\ b_{nuc} = 3.000,\ z_{nuc} = 4.200$ | 2508 | 8253 | $6.868 \times 10^1$ µm$^3$ |
| NE | | 1821 | 3548 | $4.569 \times 10^1$ µm$^2$ |
| | $r_{NP} = 0.1, h_{NP} = 1.0, p_{NP} = 1.0, R_c = 16.55$ | | | |
| Cyto | $a = 4268\ , b = 0.4244\ , c = 14.53\ , d = 23.70$ | 39660 | 184374 | $4.366 \times 10^2$ µm$^3$ |
| PM | | 16960 | 32946 | $3.431 \times 10^2$ µm$^2$ |
| Nuc | $a_{nuc} = 6.625\ , b_{nuc} = 3.000\ , z_{nuc} = 4.200$ | 2735 | 9066 | $6.870 \times 10^1$ µm$^3$ |
| NE | | 1970 | 3840 | $4.570 \times 10^1$ µm$^2$ |
| | $r_{NP} = 0.5, h_{NP} = 1.0, p_{NP} = 5.0, R_c = 20.01$ | | | |
| Cyto | $a = 2815\ , b = 0.4390\ , c = 23.66\ , d = 11.38$ | 14965 | 67920 | $4.367 \times 10^2$ µm$^3$ |
| PM | | 4596 | 8789 | $3.560 \times 10^2$ µm$^2$ |
| Nuc | $a_{nuc} = 7.427\ , b_{nuc} = 2.387\ , z_{nuc} = 3.587$ | 2652 | 8928 | $6.858 \times 10^1$ µm$^3$ |
| NE | | 1923 | 3748 | $5.450 \times 10^1$ µm$^2$ |
| | $r_{NP} = 0.25, h_{NP} = 1.0, p_{NP} = 2.5, R_c = 17.45$ | | | |
| | 0.0 µm nuclear shift | | | |
| Cyto | $a = 3383\ , b = 0.4265\ , c = 17.13\ , d = 20.07$ | 24830 | 116463 | $4.362 \times 10^2$ µm$^3$ |
| PM | | 8234 | 15928 | $3.065 \times 10^2$ µm$^2$ |
| Nuc | $a_{nuc} = 6.625\ , b_{nuc} = 3.000\ , z_{nuc} = 4.200$ | 2559 | 8439 | $6.869 \times 10^1$ µm$^3$ |
| NE | | 1861 | 3626 | $4.570 \times 10^1$ µm$^2$ |
| | 0.6 µm nuclear shift | | | |
| Cyto | $a = 3383\ , b = 0.4265\ , c = 17.13\ , d = 20.07$ | 24851 | 116920 | $4.362 \times 10^2$ µm$^3$ |
| PM | | 8233 | 15926 | $3.065 \times 10^2$ µm$^2$ |
| Nuc | $a_{nuc} = 6.625\ , b_{nuc} = 3.000\ , z_{nuc} = 4.800$ | 2241 | 7136 | $6.864 \times 10^1$ µm$^3$ |
| NE | | 1687 | 3287 | $4.567 \times 10^1$ µm$^2$ |
| | 1.0 µm nuclear shift | | | |
| Cyto | $a = 3383\ , b = 0.4265\ , c = 17.13\ , d = 20.07$ | 24902 | 117519 | $4.362 \times 10^2$ µm$^3$ |
| PM | | 8234 | 15928 | $3.065 \times 10^2$ µm$^2$ |
| Nuc | $a_{nuc} = 6.625\ , b_{nuc} = 3.000\ , z_{nuc} = 5.200$ | 2086 | 6554 | $6.864 \times 10^1$ µm$^3$ |
| NE | | 1601 | 3120 | $4.567 \times 10^1$ µm$^2$ |
| | 1.4 µm nuclear shift | | | |
| Cyto | $a = 3383\ , b = 0.4265\ , c = 17.13\ , d = 20.07$ | 24925 | 117745 | $4.362 \times 10^2$ µm$^3$ |
| PM | | 8234 | 15928 | $3.065 \times 10^2$ µm$^2$ |
| Nuc | $a_{nuc} = 6.626\ , b_{nuc} = 2.980\ , z_{nuc} = 5.600$ | 1983 | 6246 | $6.866 \times 10^1$ µm$^3$ |
| NE | | 1527 | 2975 | $4.578 \times 10^1$ µm$^2$ |

Continued on next page

| | Constants (in μm) | Vertices | Cells | Size |
|---|---|---|---|---|
| | flat, $E_{mod} = 10$ GPa, $R_c = 22.48$ | | | |
| Cyto | $a = 2590$ , $b = 0.4059$ , $c = 26.64$ , $d = 7.967$ | 4499 | 17920 | $4.355 \times 10^2$ μm$^3$ |
| PM | | 2298 | 4404 | $4.169 \times 10^2$ μm$^2$ |
| Nuc | $a_{nuc} = 6.625$ , $b_{nuc} = 3.000$ , $z_{nuc} = 3.200$ | 550 | 1846 | $6.868 \times 10^1$ μm$^3$ |
| NE | | 276 | 486 | $4.571 \times 10^1$ μm$^2$ |
| | $r_{NP} = 0.5, h_{NP} = 3.0, p_{NP} = 3.5, R_c = 15.5$ | | | |
| | $\Delta z_{indent} = 0$ μm | | | |
| Cyto | $a = 6110$ , $b = 0.5586$ , $c = 13.23$ , $d = 15.62$ | 28695 | 136211 | $4.391 \times 10^2$ μm$^3$ |
| PM | | 9188 | 17770 | $3.111 \times 10^2$ μm$^2$ |
| Nuc | $a_{nuc} = 7.179$ , $b_{nuc} = 2.555$ , $z_{nuc} = 5.755$ | 2749 | 9340 | $6.862 \times 10^1$ μm$^3$ |
| NE | | 1943 | 3786 | $4.958 \times 10^1$ μm$^2$ |
| | $\Delta z_{indent} = 0.4$ μm | | | |
| Cyto | $a = 6110$ , $b = 0.5586$ , $c = 13.23$ , $d = 15.62$ | 28585 | 135643 | $4.390 \times 10^2$ μm$^3$ |
| PM | | 9191 | 17776 | $3.111 \times 10^2$ μm$^2$ |
| Nuc | $a_{nuc} = 6.920$ , $b_{nuc} = 2.755$ , $z_{nuc} = 5.555$ | 2918 | 10026 | $6.868 \times 10^1$ μm$^3$ |
| NE | | 2029 | 3953 | $4.781 \times 10^1$ μm$^2$ |
| | $\Delta z_{indent} = 0.8$ μm | | | |
| Cyto | $a = 6110$ , $b = 0.5586$ , $c = 13.23$ , $d = 15.62$ | 28507 | 134895 | $4.390 \times 10^2$ μm$^3$ |
| PM | | 9185 | 17764 | $3.111 \times 10^2$ μm$^2$ |
| Nuc | $a_{nuc} = 6.701$ , $b_{nuc} = 2.955$ , $z_{nuc} = 5.355$ | 3174 | 11099 | $6.871 \times 10^1$ μm$^3$ |
| NE | | 2149 | 4187 | $4.691 \times 10^1$ μm$^2$ |
| | $\Delta z_{indent} = 1.0$ μm | | | |
| Cyto | $a = 6110$ , $b = 0.5586$ , $c = 13.23$ , $d = 15.62$ | 28491 | 134564 | $4.390 \times 10^2$ μm$^3$ |
| PM | | 9185 | 17764 | $3.111 \times 10^2$ μm$^2$ |
| Nuc | $a_{nuc} = 6.663$ , $b_{nuc} = 3.000$ , $z_{nuc} = 5.200$ | 3302 | 11597 | $6.868 \times 10^1$ μm$^3$ |
| NE | | 2238 | 4362 | $4.717 \times 10^1$ μm$^2$ |
| | $\Delta z_{indent} = 1.4$ μm | | | |
| Cyto | $a = 6110$ , $b = 0.5586$ , $c = 13.23$ , $d = 15.62$ | 28367 | 133232 | $4.391 \times 10^2$ μm$^3$ |
| PM | | 9182 | 17758 | $3.111 \times 10^2$ μm$^2$ |
| Nuc | $a_{nuc} = 6.698$ , $b_{nuc} = 3.000$ , $z_{nuc} = 4.800$ | 3617 | 12913 | $6.862 \times 10^1$ μm$^3$ |
| NE | | 2414 | 4703 | $4.886 \times 10^1$ μm$^2$ |
| | $\Delta z_{indent} = 1.8$ μm | | | |
| Cyto | $a = 6110$ , $b = 0.5586$ , $c = 13.23$ , $d = 15.62$ | 28322 | 132753 | $4.392 \times 10^2$ μm$^3$ |
| PM | | 9173 | 17740 | $3.111 \times 10^2$ μm$^2$ |
| Nuc | $a_{nuc} = 6.733$ , $b_{nuc} = 3.000$ , $z_{nuc} = 4.400$ | 3995 | 14456 | $6.852 \times 10^1$ μm$^3$ |
| NE | | 2640 | 5143 | $5.102 \times 10^1$ μm$^2$ |
| | $\Delta z_{indent} = 2.0$ μm | | | |
| Cyto | $a = 6110$ , $b = 0.5586$ , $c = 13.23$ , $d = 15.62$ | 28407 | 132884 | $4.393 \times 10^2$ μm$^3$ |
| PM | | 9182 | 17758 | $3.111 \times 10^2$ μm$^2$ |
| Nuc | $a_{nuc} = 6.749$ , $b_{nuc} = 3.000$ , $z_{nuc} = 4.200$ | 4210 | 15284 | $6.839 \times 10^1$ μm$^3$ |
| NE | | 2776 | 5408 | $5.219 \times 10^1$ μm$^2$ |

Continued on next page

| | Constants (in µm) | Vertices | Cells | Size |
|---|---|---|---|---|
| | $\Delta z_{indent} = 2.4$ µm | | | |
| Cyto | $a = 6110$ , $b = 0.5586$ , $c = 13.23$ , $d = 15.62$ | 28267 | 131547 | $4.396 \times 10^2$ µm$^3$ |
| PM | | 9172 | 17738 | $3.111 \times 10^2$ µm$^2$ |
| Nuc | $a_{nuc} = 6.779$ , $b_{nuc} = 3.000$ , $z_{nuc} = 3.800$ | 4538 | 16656 | $6.811 \times 10^1$ µm$^3$ |
| NE | | 2969 | 5781 | $5.463 \times 10^1$ µm$^2$ |
| | $\Delta z_{indent} = 2.8$ µm | | | |
| Cyto | $a = 6110$ , $b = 0.5586$ , $c = 13.23$ , $d = 15.62$ | 28216 | 130846 | $4.400 \times 10^2$ µm$^3$ |
| PM | | 9160 | 17716 | $3.111 \times 10^2$ µm$^2$ |
| Nuc | $a_{nuc} = 6.805$ , $b_{nuc} = 3.000$ , $z_{nuc} = 3.400$ | 4869 | 17912 | $6.768 \times 10^1$ µm$^3$ |
| NE | | 3193 | 6215 | $5.710 \times 10^1$ µm$^2$ |

### S2 Equations and parameters for YAP/TAZ mechanotransduction model

The model is summarized in terms of species, reactions, and parameters according to the SMART framework [6]. For clarity, the full system of PDEs is given explicitly at the end of this section.

**Table S2:** Species in YAP/TAZ mechanotransduction model. All initial conditions and diffusion coefficients are used from [2], except YAP/TAZ diffusion coefficients from [7].

| | Compartment | $D$ (µm$^2$ s$^{-1}$) | Initial Condition |
|---|---|---|---|
| Emod* | PM | 0 | 10 GPa |
| pFAK | Cyto | 10 | $3.000 \times 10^{-1}$ µM |
| RhoA_GDP | Cyto | 1 | 1.000 µM |
| RhoA_GTP | PM | 0.3 | $3.360 \times 10^1$ µm$^{-2}$ |
| ROCK_A | Cyto | 75 | 0.000 µM |
| mDia_A | Cyto | 1 | $1.000 \times 10^{-2}$ µM |
| Myo_A | Cyto | 0.8 | 1.500 µM |
| LIMK_A | Cyto | 10 | $1.000 \times 10^{-1}$ µM |
| Cofilin_NP | Cyto | 10 | 1.800 µM |
| FActin | Cyto | 0.6 | $1.790 \times 10^1$ µM |
| GActin | Cyto | 13.37 | $4.824 \times 10^2$ µM |
| LaminA | NE | 0.001 | 0.000 µm$^{-2}$ |
| NPC_A | NE | 0.001 | 0.000 µm$^{-2}$ |
| YAPTAZ_free | Cyto | 80.0 | $7.000 \times 10^{-1}$ µM |
| YAPTAZ_seq | Cyto | 80.0 | $2.000 \times 10^{-1}$ µM |
| YAPTAZ_nuc | Nuc | 4.0 | $7.000 \times 10^{-1}$ µM |

*Emod represents the substrate stiffness and also specifies the location of the substrate ($z = 0$ or surface of nanopillars).

**Table S3:** Reactions in YAP/TAZ mechanotransduction model.

| | Reactants | Products | Equation |
|---|---|---|---|
| $a_1$ | [] | ['pFAK'] | $cyto_{convert}([FAK_{tot}] - [pFAK]) \left(\frac{\rho_{I,bound}}{\rho_{I,max}}\right)\left(\frac{E_{mod}}{C + E_{mod}} k_{sf} + k_f\right)$ where $\left(\frac{\rho_{I,bound}}{\rho_{I,max}}\right) = \exp\left(-\frac{H_{PM}}{H_0}\right)\mathcal{H}(H_{PM}) + \mathcal{H}(-H_{PM})$ on $\Gamma_{substrate}$, $\mathcal{H}$ is the Heaviside function, and $\Gamma_{substrate}$ is the PM in contact with the substrate. |

Continued on next page

| | Reactants | Products | Equation |
|---|---|---|---|
| $a_2$ | [‘pFAK’] | [] | $k_{df}[pFAK]$ |
| $a_3$ | [‘RhoA_GDP’] | [‘RhoA_GTP’] | $[RhoA_{GDP}]cyto_{convert}k_{fkrho}(\gamma[pFAK]^5+1) - \frac{[RhoA_{GTP}]k_{drho}}{N_{conv}}$ |
| $a_4$ | [] | [‘ROCK_A’] | $[RhoA_{GTP}]k_{rrho}([ROCK_{tot}]-[ROCK_A])$ |
| $a_5$ | [‘ROCK_A’] | [] | $[ROCK_A]k_{drock}$ |
| $b_1$ | [] | [‘mDia_A’] | $[RhoA_{GTP}]k_{mrho}([mDia_{tot}]-[mDia_A])$ |
| $b_2$ | [‘mDia_A’] | [] | $k_{dmdia}[mDia_A]$ |
| $b_3$ | [] | [‘Myo_A’] | $k_{mr}([Myo_{tot}]-[Myo_A])\cdot \left([ROCK_A]\epsilon\frac{1+\tanh(sc1([ROCK_A]-[ROCK_B]))}{2}+1\right) -[Myo_A]k_{dmy}$ |
| $b_4$ | [] | [‘LIMK_A’] | $k_{lr}([LIMK_{tot}]-[LIMK_A])\cdot \left([ROCK_A]\tau\frac{1+\tanh(sc1([ROCK_A]-[ROCK_B]))}{2}+1\right) -[LIMK_A]k_{dl}$ |
| $b_5$ | [] | [‘Cofilin_NP’] | $k_{turnover}([Cofilin_{tot}]-[Cofilin_{NP}]) - \frac{[Cofilin_{NP}][LIMK_A]k_{catCof}}{Cofilin_{NP}+k_{mCof}}$ |
| $b_6$ | [‘GActin’] | [‘FActin’] | $[GActin]k_{ra}\cdot \left(\alpha_m[mDia_A]\frac{1+\tanh(sc1([mDia_A]-[mDia_B]))}{2}+1\right) -[FActin]([Cofilin_{NP}]k_{fc1}+k_{dep})$ |
| $b_7$ | [‘GActin’] | [‘FActin’] | $k_{N\text{-}WASP}\exp\left(\frac{H_{PM}}{H_{N\text{-}WASP}}\right)[GActin]$ |
| $c_1$ | [‘YAPTAZ_seq’] | [‘YAPTAZ_free’] | $[YAPTAZ_{seq}]([FActin][Myo_A]k_{CY}+k_{CN}) -[YAPTAZ_{free}]k_{NC}$ |
| $c_3$ | [] | [‘LaminA’] | $\frac{[FActin]^{2.6}k_{fl}p([LaminA_{tot}]-[LaminA])}{C_{LaminA}+[FActin]^{2.6}p} - [LaminA]k_{rl}$ |
| $c_4$ | [] | [‘NPC_A’] | $[FActin][LaminA][Myo_A]k_{fNPC}([NPC_{tot}]-[NPC_A]) -[NPC_A]k_{rNPC}$ |

Continued on next page

| | Reactants | Products | Equation |
|---|---|---|---|
| $c_5$ | ['YAPTAZ_free'] | ['YAPTAZ_nuc'] | $[YAPTAZ_{free}]([NPC_A]k_{in} + k_{inb}) \exp\left(\frac{\alpha - 1}{\alpha_0}\right)$ $-[YAPTAZ_{nuc}]k_{out}$ |
| $c_6$ | ['YAPTAZ_free'] | ['YAPTAZ_nuc'] | $p_{NER}k_{rupture}([YAPTAZ_{free}] - \frac{[YAPTAZ_{nuc}]}{\upsilon})$ |

**Table S4:** Parameters in YAP/TAZ mechanotransduction model. Values were adopted directly from those reported in [2] unless otherwise noted.

| | Value | Description |
|---|---|---|
| $FAK_{tot}$ | $1.000\,\mu M$ | Total cytosolic FAK |
| $k_f$ | $1.500 \times 10^{-2}\,s^{-1}$ | rate constant for baseline FAK phosphorylation |
| $k_{sf}$ | $3.790 \times 10^{-1}\,s^{-1}$ | substrate-stiffness-dependent FAK phosphorylation rate constant |
| $E_{mod}$ | $1 \times 10^{7}\,kPa$ | substrate stiffness |
| $C$ | $3.250\,kPa$ | critical substrate stiffness |
| $cyto_{convert}$ | $1.825\,\mu m$ | vol/SA ratio |
| $N_{conv}$ | $6.022 \times 10^{2}\,\mu m^{-3}\,\mu M^{-1}$ | unit conversation factor |
| $H_{PM}$ | Equation (S36) ($\mu m^{-1}$) | plasma membrane mean curvature |
| $H_{0,FAK}$ | varied ($\mu m^{-1}$) | curvature sensitivity of FAK phosphorylation |
| $k_{df}$ | $3.500 \times 10^{-2}\,s^{-1}$ | FAK dephosphorylation rate constant |
| $k_{fkrho}$ | $1.680 \times 10^{-2}\,s^{-1}$ | RhoA activation rate constant |
| $\gamma$ | $7.756 \times 10^{1}\,\mu M^{-5}$ | Scaling factor |
| $k_{drho}$ | $6.250 \times 10^{-1}\,s^{-1}$ | RhoA deactivation rate constant |
| $k_{rrho}$ | $6.480 \times 10^{-1}\,\mu M^{-1}\,s^{-1}$ | ROCK activation rate constant |
| $ROCK_{tot}$ | $1.000\,\mu M$ | Total cytosolic ROCK |
| $k_{drock}$ | $8.000 \times 10^{-1}\,s^{-1}$ | ROCK deactivation rate constant |
| $k_{mrho}$ | $2.000 \times 10^{-3}\,\mu M^{-1}\,s^{-1}$ | mDia activation rate constant |
| $mDia_{tot}$ | $8.000 \times 10^{-1}\,\mu M$ | Total cytosolic mDia |
| $k_{dmdia}$ | $5.000 \times 10^{-3}\,s^{-1}$ | mDia deactivation rate constant |
| $Myo_{tot}$ | $5.000\,\mu M$ | Total cytosolic myosin |
| $k_{mr}$ | $3.000 \times 10^{-2}\,s^{-1}$ | Myosin activation rate constant |
| $ROCK_B$ | $3.000 \times 10^{-1}\,\mu M$ | Critical activated ROCK concentration |
| $\epsilon$ | $3.600 \times 10^{1}\,\mu M^{-1}$ | Scaling factor |
| $sc1$ | $2.000 \times 10^{1}\,\mu M^{-1}$ | Scaling factor |
| $k_{dmy}$ | $6.700 \times 10^{-2}\,s^{-1}$ | Myosin deactivation rate constant |
| $LIMK_{tot}$ | $2.000\,\mu M$ | Total cytosolic LIMK |
| $k_{lr}$ | $7.000 \times 10^{-2}\,s^{-1}$ | LIMK activation rate constant |
| $\tau$ | $5.549 \times 10^{1}\,\mu M^{-1}$ | Scaling factor |
| $k_{dl}$ | $2.000\,s^{-1}$ | LIMK deactivation rate constant |
| $Cofilin_{tot}$ | $2.000\,\mu M$ | Total cytosolic cofilin |
| $k_{turnover}$ | $4.000 \times 10^{-2}\,s^{-1}$ | Cofilin dephosphorylation rate constant |
| $k_{catCof}$ | $3.400 \times 10^{-1}\,s^{-1}$ | Cofilin phosphorylation rate constant |
| $k_{mCof}$ | $4.000\,\mu M$ | Critical cofilin concentration |
| $k_{ra}$ | $4.000 \times 10^{-1}\,s^{-1}$ | Actin polymerization rate constant |
| $\alpha_m$ | $5.000 \times 10^{1}\,\mu M^{-1}$ | Scaling factor |
| $mDia_B$ | $1.650 \times 10^{-1}\,\mu M$ | Critical activated mDia concentration |
| $k_{dep}$ | $3.500\,s^{-1}$ | Actin depolymerization rate constant |
| $k_{fc1}$ | $4.000\,\mu M^{-1}\,s^{-1}$ | Cofilin-mediated actin depolymerization rate constant |
| $k_{N\text{-}WASP}$ | $1.000 \times 10^{-2}\,\mu m\,s^{-1}$ | N-WASP-mediated actin polymerization rate constant (see main text) |

Continued on next page

| | Value | Description |
|---|---|---|
| $H_{N\text{-}WASP}$ | $2.000\,\mu m^{-1}$ | N-WASP curvature sensitivity (see main text and [8, 9]) |
| $k_{CN}$ | $5.600 \times 10^{-1}\,s^{-1}$ | Baseline YAP/TAZ desequestration rate constant |
| $k_{CY}$ | $7.600 \times 10^{-4}\,\mu M^{-2}\,s^{-1}$ | stress-fiber-mediated YAP/TAZ de-sequestration rate constant |
| $k_{NC}$ | $1.400 \times 10^{-1}\,s^{-1}$ | YAP/TAZ sequestration rate constant |
| $LaminA_{tot}$ | $3.500 \times 10^{3}\,\mu m^{-2}$ | Total lamin A in NE |
| $k_{fl}$ | $4.600 \times 10^{-1}\,s^{-1}$ | Lamin A dephosphorylation rate constant |
| $p$ | $9.000 \times 10^{-6}\,kPa\,\mu M^{-2.6}$ | Scaling factor for cell stiffness |
| $C_{LaminA}$ | $1.000 \times 10^{2}\,kPa$ | Critical cell stiffness for lamin A dephosphorylation |
| $k_{rl}$ | $1.000 \times 10^{-3}\,s^{-1}$ | Lamin A phosphorylation rate constant |
| $NPC_{tot}$ | $6.500\,\mu m^{-2}$ | Total NPC density in NE |
| $k_{fNPC}$ | $2.800 \times 10^{-7}\,\mu m^{2}\,\mu M^{-2}\,s^{-1}$ | NPC opening rate constant |
| $k_{rNPC}$ | $8.700\,s^{-1}$ | NPC closing rate constant |
| $k_{inb}$ | $1.000\,\mu m^{-2}\,\mu M^{-1}\,s^{-1}$ | NPC-independent YAP/TAZ nuclear import rate |
| $k_{in}$ | $1.000 \times 10^{1}\,\mu M^{-1}\,s^{-1}$ | NPC-dependent YAP/TAZ nuclear import rate |
| $k_{out}$ | $1.000\,\mu m^{-2}\,\mu M^{-1}\,s^{-1}$ | YAP/TAZ nuclear export rate |
| $\alpha_{NE}$ | Equation (S47) (unitless) | stretch ratio of NE |
| $\alpha_{NE,0}$ | varied (unitless) | NPC sensitivity to stretch |
| $\upsilon$ | varied (unitless) | ratio of import to export through rupture-induced pore |
| $k_{rupture}$ * | $1.000 \times 10^{5}\,s^{-1}\,\mu M^{-1}\,\mu m^{-2}$ | rate of export through rupture-induced pore |
| $p_{NER}$ | $\left(1 - \exp\left(-\frac{t}{t_0}\right)\right) \cdot \exp\left(-\frac{x^2+y^2+(z-z_{NE})^2}{\sigma_{pore}^2}\right)$ | relative permeability of rupture-induced pore |
| $t_0$ | $1.000\,s$ | time constant for pore formation |
| $z_{NE}$ | $h_{NP} + 0.2\,\mu m$ | z-value of NE at which rupture-induced pore forms |
| $\sigma_{pore}$ | varied (μm) | effective radius of rupture-induced pore |

* Order of magnitude for $k_{rupture}$ is estimated assuming diffusive transport; i.e., $k_{rupture} = N_{convert}\frac{D_{Y,NER}}{d_{NE}}$, where $D_{Y,NER}$ is the effective diffusion coefficient for transport through the NER region (assumed to be between YAP/TAZ diffusion coefficients in the cytosol and nucleus, at about 10 μm $^2$/s) and $d_{NE}$ is the thickness of the NE (about 50 nm).

The best-fit value of $H_0$ was determined by assessing the normalized sum of square errors ($NSSE$) between experimental data from [10] and our model over the 5 nanopillar substrates considered here:

$$NSSE = \sum_{k=1}^{5} \left(\frac{1}{SEM_{exp,k}}\right)^2 \left[\left(\frac{Y_{nuc}}{Y_{cyto}}\right)_{exp,k} - \left(\frac{Y_{nuc}}{Y_{cyto}}\right)_{sim,k}\right]^2, \qquad \text{(S9)}$$

where $SEM_{exp,k}$ is the standard error of the mean associated with the YAP/TAZ N/C measured on substrate $k$, and $\left(\frac{Y_{nuc}}{Y_{cyto}}\right)_{exp,k}$ and $\left(\frac{Y_{nuc}}{Y_{cyto}}\right)_{sim,k}$ are the mean experimental YAP/TAZ N/C and the spatially averaged YAP/TAZ N/C for substrate $k$, respectively. The $NSSE$ is plotted as a function of $H_0$ for two different values of $k_{N\text{-}WASP}$ in Fig S1B.

Given the equations, parameters, and variables defined in the above equations, we can write out the full system of partial differential equations that are solved in SMART. Volume domains are written as $\Omega$ (i.e., $\Omega_{cyto}$ and $\Omega_{nuc}$) and surface domains are written as $\Gamma$ (i.e., $\Gamma_{PM}$ and $\Omega_{NE}$). $\nabla_S^2$ denotes the surface Laplacian operator.

$$\begin{aligned} \frac{\partial[pFAK]}{\partial t} &= D_{FAK}\nabla^2[pFAK] - a_2 \quad \text{in } \Omega_{cyto} \\ -D_{FAK}\mathbf{n}_{PM} \cdot \nabla[pFAK] &= a_1 \quad \text{on } \Gamma_{PM} \\ D_{FAK}\mathbf{n}_{NE} \cdot \nabla[pFAK] &= 0 \quad \text{on } \Gamma_{NE} \end{aligned} \qquad \text{(S10)}$$

$$\frac{\partial[RhoA_{GTP}]}{\partial t} = D_{RhoA_{GTP}} \nabla_S^2 [RhoA_{GTP}] + N_{conv} a_3 \quad \text{on } \Gamma_{PM} \tag{S11}$$

$$\begin{aligned}
\frac{\partial[RhoA_{GDP}]}{\partial t} &= D_{RhoA_{GDP}} \nabla^2 [RhoA_{GDP}] \quad \text{in } \Omega_{cyto} \\
-D_{RhoA_{GDP}} \mathbf{n}_{PM} \cdot \nabla [RhoA_{GDP}] &= -a_3 \quad \text{on } \Gamma_{PM} \\
D_{RhoA_{GDP}} \mathbf{n}_{NE} \cdot \nabla [RhoA_{GDP}] &= 0 \quad \text{on } \Gamma_{NE}
\end{aligned} \tag{S12}$$

$$\begin{aligned}
\frac{\partial[ROCK_A]}{\partial t} &= D_{ROCK} \nabla^2 [ROCK_A] - a_5 \quad \text{in } \Omega_{cyto} \\
-D_{ROCK} \mathbf{n}_{PM} \cdot \nabla [ROCK_A] &= a_4 \quad \text{on } \Gamma_{PM} \\
D_{ROCK} \mathbf{n}_{NE} \cdot \nabla [ROCK_A] &= 0 \quad \text{on } \Gamma_{NE}
\end{aligned} \tag{S13}$$

$$\begin{aligned}
\frac{\partial[mDia_A]}{\partial t} &= D_{mDia} \nabla^2 [mDia_A] - b_2 \quad \text{in } \Omega_{cyto} \\
-D_{mDia} \mathbf{n}_{PM} \cdot \nabla [mDia_A] &= b_1 \quad \text{on } \Gamma_{PM} \\
D_{mDia} \mathbf{n}_{NE} \cdot \nabla [mDia_A] &= 0 \quad \text{on } \Gamma_{NE}
\end{aligned} \tag{S14}$$

$$\begin{aligned}
\frac{\partial[Myo_A]}{\partial t} &= D_{Myo} \nabla^2 [Myo_A] + b_3 \quad \text{in } \Omega_{cyto} \\
-D_{Myo} \mathbf{n}_{PM} \cdot \nabla [Myo_A] &= 0 \quad \text{on } \Gamma_{PM} \\
D_{Myo} \mathbf{n}_{NE} \cdot \nabla [Myo_A] &= 0 \quad \text{on } \Gamma_{NE}
\end{aligned} \tag{S15}$$

$$\begin{aligned}
\frac{\partial[LIMK_A]}{\partial t} &= D_{LIMK} \nabla^2 [LIMK_A] + b_4 \quad \text{in } \Omega_{cyto} \\
-D_{LIMK} \mathbf{n}_{PM} \cdot \nabla [LIMK_A] &= 0 \quad \text{on } \Gamma_{PM} \\
D_{LIMK} \mathbf{n}_{NE} \cdot \nabla [LIMK_A] &= 0 \quad \text{on } \Gamma_{NE}
\end{aligned} \tag{S16}$$

$$\begin{aligned}
\frac{\partial[Cofilin_{NP}]}{\partial t} &= D_{Cofilin} \nabla^2 [Cofilin_{NP}] + b_5 \quad \text{in } \Omega_{cyto} \\
-D_{Cofilin} \mathbf{n}_{PM} \cdot \nabla [Cofilin_{NP}] &= 0 \quad \text{on } \Gamma_{PM} \\
D_{Cofilin} \mathbf{n}_{NE} \cdot \nabla [Cofilin_{NP}] &= 0 \quad \text{on } \Gamma_{NE}
\end{aligned} \tag{S17}$$

$$\begin{aligned}
\frac{\partial[FActin]}{\partial t} &= D_{FActin} \nabla^2 [FActin] + b_6 \quad \text{in } \Omega_{cyto} \\
-D_{FActin} \mathbf{n}_{PM} \cdot \nabla [FActin] &= b_7 \quad \text{on } \Gamma_{PM} \\
D_{FActin} \mathbf{n}_{NE} \cdot \nabla [FActin] &= 0 \quad \text{on } \Gamma_{NE}
\end{aligned} \tag{S18}$$

(S19)

$$\begin{aligned}\frac{\partial[YAPTAZ_{seq}]}{\partial t} &= D_{YAPTAZ,cyto}\nabla^2[YAPTAZ_{seq}] - c_1 \quad \text{in } \Omega_{cyto} \\ -D_{YAPTAZ,cyto}\mathbf{n}_{PM}\cdot\nabla[YAPTAZ_{seq}] &= 0 \quad \text{on } \Gamma_{PM} \\ D_{YAPTAZ,cyto}\mathbf{n}_{NE}\cdot\nabla[YAPTAZ_{seq}] &= 0 \quad \text{on } \Gamma_{NE}\end{aligned} \tag{S20}$$

$$\begin{aligned}\frac{\partial[YAPTAZ_{free}]}{\partial t} &= D_{YAPTAZ,cyto}\nabla^2[YAPTAZ_{free}] + c_1 \quad \text{in } \Omega_{cyto} \\ -D_{YAPTAZ,cyto}\mathbf{n}_{PM}\cdot\nabla[YAPTAZ_{free}] &= 0 \quad \text{on } \Gamma_{PM} \\ D_{YAPTAZ,cyto}\mathbf{n}_{NE}\cdot\nabla[YAPTAZ_{free}] &= -\frac{c_5+c_6}{N_{conv}} \quad \text{on } \Gamma_{NE}\end{aligned} \tag{S21}$$

$$\frac{\partial[LaminA]}{\partial t} = D_{LaminA}\nabla_S^2[LaminA] + c_3 \quad \text{on } \Gamma_{NE} \tag{S22}$$

$$\frac{\partial[NPC_A]}{\partial t} = D_{LaminA}\nabla_S^2[NPC_A] + c_4 \quad \text{on } \Gamma_{NE} \tag{S23}$$

$$\begin{aligned}\frac{\partial[YAPTAZ_{nuc}]}{\partial t} &= D_{YAPTAZ,nuc}\nabla^2[YAPTAZ_{nuc}] \quad \text{in } \Omega_{cyto} \\ -D_{YAPTAZ,nuc}\mathbf{n}_{NE}\cdot\nabla[YAPTAZ_{nuc}] &= \frac{c_5+c_6}{N_{conv}} \quad \text{on } \Gamma_{NE}\end{aligned} \tag{S24}$$

### S3 Well-mixed approximation for YAP/TAZ N/C

Here, we assume that YAP/TAZ is well-mixed within each compartment (the cytosol and the nucleus), but other variables may still exhibit some spatial dependence. In this case, dynamics governing YAP/TAZ nuclear translocation is governed by two ODEs. Writing concentrations of free cytosolic YAP/TAZ, sequestered cytosolic YAP/TAZ, and nuclear YAP/TAZ as $Y_{free}$, $Y_{seq}$, and $Y_{nuc}$, respectively, the volume of the nucleus as $vol_{nuc}$, and the volume of the cytosol as $vol_{cyto}$:

$$\frac{dY_{free}}{dt} = (k_{CN} + \bar{k}_{CY})Y_{seq} - k_{NC}Y_{free} + \frac{1}{N_{conv}vol_{cyto}}\left(k_{out,tot}Y_{nuc} - k_{in,tot}Y_{free}\right) \tag{S25}$$

$$\frac{dY_{nuc}}{dt} = \frac{1}{N_{conv}vol_{nuc}}\left(k_{in,tot}Y_{free} - k_{out,tot}Y_{nuc}\right) \tag{S26}$$

$$\begin{aligned}&\text{where} \quad Y_{seq} = \frac{Y_{tot} - N_{conv}(Y_{nuc}vol_{nuc} + Y_{free}vol_{cyto})}{N_{conv}vol_{cyto}}, \\ &k_{in,tot} = \bar{k}_{in,NPC} + p_{NER}k_{rupture}SA_{NER}, \\ &\text{and} \quad k_{out,tot} = k_{out,NPC}SA_{nuc} + p_{NER}\frac{k_{rupture}}{v}SA_{NER}\end{aligned}$$

$\bar{k}_{in,NPC}$ is defined to account for spatially dependent stretch-sensitive entry:

$$\bar{k}_{in,NPC} = \int_{\Gamma_{NE}} (k_{inb} + k_{in}[NPC_A])\exp\left(\frac{\alpha-1}{\alpha_0}\right)d\Gamma, \tag{S27}$$

and $\bar{k}_{CY}$ is defined to account for actomyosin activity across the entire cell:

$$\bar{k}_{CY} = \frac{1}{vol_{cyto}} \int_{\Omega_{cyto}} k_{CY}[FActin][Myo_A]d\Omega. \tag{S28}$$

Assuming the rupture-induced pore is located over a flat patch of membrane directly above the central nanopillar, the effective surface area occupied by the pore upon completion of its formation is:

$$SA_{NER} = \int_{-\infty}^{\infty}\int_{-\infty}^{\infty} \exp\left(-\frac{x^2+y^2}{\sigma_{pore}^2}\right) dxdy = \pi\sigma_{pore}^2. \tag{S29}$$

The resulting steady-state YAP/TAZ N/C is given by:

$$\begin{aligned} \frac{Y_{nuc}}{Y_{free}+Y_{seq}} &= \frac{\phi_{free}(\bar{k}_{in,NPC}+k_{rupture}SA_{NER})}{k_{out,NPC}SA_{nuc}+\frac{k_{rupture}}{\upsilon}SA_{NER}}, \\ \text{where} \quad \phi_{free} &= \frac{k_{CN}+\bar{k}_{CY}}{k_{NC}+k_{CN}+\bar{k}_{CY}}. \end{aligned} \tag{S30}$$

We consider a couple specialized cases presented in the main text. In these examples, the NE is split into two regions - one uniformly stretched with associated area dilation $\alpha$ and activated NPC density $[NPC_A]_{def}$ and the other remaining undeformed ($\alpha = 1$ and $[NPC_A] = [NPC_A]_{undef}$). This differs from the non-uniform stretch introduced in Section S5, but introduces a simplified form of Equation (S27):

$$\bar{k}_{in,NPC} = (k_{inb} + k_{in}[NPC_A]_{def}) \exp\left(\frac{\alpha-1}{\alpha_0}\right) SA_{def} + (k_{inb} + k_{in}[NPC_A]_{undef}) SA_{undef}, \tag{S31}$$

When $k_{rupture} = 0$ (no NER), Equation (S30) then simplifies to the form in the main text (Eq 5). For small deformations over a region of the reference geometry with surface area $SA_0$, $SA_{def} \approx \alpha SA_0$. For illustration, we plot the predictions of Equation (S31) as a function of $\alpha$ (all other parameters fixed) in Figure S3B.

When $k_{rupture} \neq 0$, we obtain the remaining case in Eq 10 of the main text. For illustration, we then plot the predictions of Equation (S31) as a function of $k_{rupture}SA_{NER}$ (all other parameters fixed) in Figure S3C.

### S4 Plasma membrane curvature calculations

Here, we describe our strategy to define plasma membrane shape around nanopillars. Rather than derive membrane morphology from first principles (e.g., energy minimization), we describe cell shape using empirical relationships that match well with cell morphology in experimental images (cf. Fig 3). The curvature of the plasma membrane can then be analytically expressed in the region surrounding each nanopillar.

The membrane shape around each nanopillar is assumed to be axisymmetric with respect to the central axis of the pillar, with a cross-sectional profile composed of lines and circular arcs (Figure S4A-B) We assume that the membrane is attached to the nanopillar surface via adhesive interactions, fixed at a minimum distance of $d_{steric} =$ 50 nm from the surface due to steric repulsion. Furthermore, the membrane curves to meet the lower surface with a characteristic radius, $d_{curv} = 200$ nm, as depicted in Figure S4A.

The segments of this geometry are readily parameterized in terms of arc length $s$ in the $r_{local}, z$ plane, where $r_{local}$ is the radial coordinate defined with the origin at the center of a given nanopillar located at the point $(x_{NP}, y_{NP})$. That is, $r_{local} = \sqrt{(x-x_{NP})^2 + (y-y_{NP})^2}$, where $x$ and $y$ are the global Cartesian coordinates (all coordinates defined in Figure S4A-B). For a nanopillar of radius $r_{NP}$ and height $h_{NP}$:

$$r_{local} = \begin{cases} s & 0 \le s < s_1 \\ r_{NP} + d_{steric} \sin\phi & s_1 \le s < s_2 \\ r_{NP} + d_{steric} & s_2 \le s < s_3 \\ r_{NP} + d_{steric} + d_{curv}(1 - \sin\phi) & s_3 \le s < s_4 \\ r_{NP} + d_{steric} + d_{curv} + s - s_4 & s \ge s_4 \end{cases}, \tag{S32}$$

and

$$z = \begin{cases} h_{NP} & 0 \le s < s_1 \\ h_{NP} + d_{steric}(\cos\phi - 1) & s_1 \le s < s_2 \\ h_{NP} - d_{steric} - (s - s_2) & s_2 \le s < s_3 \\ d_{curv}(1 - \cos\phi) & s_3 \le s < s_4 \\ 0 & s \ge s_4 \end{cases}, \tag{S33}$$

The arc length values $s_1$, $s_2$, $s_3$, and $s_4$ are shown in Figure S4B and can be expressed in terms of the defined geometric parameters:

$$\begin{aligned} s_1 &= r_{NP} \\ s_2 &= s_1 + \frac{\pi d_{steric}}{2} \\ s_3 &= s_2 + h_{NP} - d_{steric} - d_{curv} \\ s_4 &= s_3 + \frac{\pi d_{curv}}{2} \end{aligned} \tag{S34}$$

$\phi$ is the polar angle measured between the normal vector $\mathbf{n}$ (Fig S4A) and the $z$-axis:

$$\phi = \begin{cases} 0 & 0 \le s < s_1 \\ \frac{s - s_1}{d_{steric}} & s_1 \le s < s_2 \\ \frac{\pi}{2} & s_2 \le s < s_3 \\ \frac{s_4 - s}{d_{curv}} & s_3 \le s < s_4 \\ 0 & s \ge s_4 \end{cases}, \tag{S35}$$

Given the axisymmetry of the geometry, curvature can be readily calculated using the following relationship:

$$H = \frac{1}{2}\left(\frac{\partial\phi}{\partial s} + \frac{\sin\phi}{r_{local}}\right) = \begin{cases} 0 & 0 \le s < s_1 \\ \frac{1}{2d_{steric}}\left(1 + \frac{r_{local} - r_{NP}}{r_{local}}\right) & s_1 \le s < s_2 \\ \frac{1}{2(r_{NP} + d_{steric})} & s_2 \le s < s_3 \\ -\frac{1}{2d_{curv}}\left(1 + \frac{r_{local} - r_{NP} - d_{curv} - d_{steric}}{r_{local}}\right) & s_3 \le s < s_4 \\ 0 & s \ge s_4 \end{cases}. \tag{S36}$$

Note that the orientation of the normal vector $\mathbf{n}$ is defined such that the sign of membrane curvature matches the convention in literature; that is, an inwardly curved membrane has positive mean curvature.

Plasma membrane curvature computed using Equation (S36) and the resulting FAK phosphorylation rate dictated by Equation 2 in the main text are shown in Fig S4C.

#### S5 Analysis of prescribed nuclear deformations

We now consider a prescribed deformation field on the nucleus. As in the case of the plasma membrane above, we formulate empirical relationships that produce shapes matching well with experiments (cf. Fig 3) rather than derive these shapes using mechanical arguments. For simplicity, we assume the nucleus adopts a reference geometry of an oblate spheroid, dictated by Equation (S5):

$$\frac{r^2}{a_{nuc}^2} + \frac{(Z - z_{nuc})^2}{b_{nuc}^2} = 1. \tag{S37}$$

For clarity, we write the reference (i.e., non-deformed) $z$-coordinate as $Z$. We assume that all deformations occur in the $z$-direction, so reference and deformed coordinates are the same in the $x$ and $y$ (and therefore, $r$) directions (Figure S5A). We allow a gap between the plasma membrane and the nuclear envelope, $\Delta z_{PM,NE}$, fixed 0.2 µm here. The $z$-component of the nuclear deformation induced by a nanopillar of radius $r_{NP}$ and height $h_{NP}$ located at $(x_{NP}, y_{NP})$ is then:

$$u_z(x,y,Z) = \begin{cases} \Delta Z_{NP} & \Delta r_{NP} \le 0 \quad \text{and} \quad \Delta Z_{NP} \ge 0 \text{ (directly above nanopillar)} \\ \Delta Z_{NP} \exp\left(-\frac{\Delta r_{NP}^2}{\sigma^2}\right) & \Delta r_{NP} > 0 \quad \text{and} \quad \Delta Z_{NP} \ge 0 \text{ (between nanopillars)} \\ 0 & \Delta Z_{NP} < 0 \text{ (upper half of the nucleus)} \end{cases}, \tag{S38}$$

where $r_{local} = \sqrt{(x - x_{NP})^2 + (y - y_{NP})^2}$, $\Delta Z_{NP} = h_{NP} + \Delta z_{PM,NE} - Z$, and $\Delta r_{NP} = r_{local} - (r_{NP} + 2d_{steric})$. For this deformation field, the nucleus is locally flat directly above the nanopillar and within $2d_{steric}$ of the edge of the nanopillar and relaxes with a Gaussian profile away from the nanopillar as shown in Figure S5B. $\sigma$ determines the width of the Gaussian profile and is fixed to 0.2 µm here. All coordinates are labeled in Figure S5C.

The local strain is characterized by the deformation gradient tensor, which is expressed in matrix form in this case as:

$$\mathbf{F} = \begin{bmatrix} 1 & 0 & 0 \\ 0 & 1 & 0 \\ u_{z,x} & u_{z,y} & 1 + u_{z,Z} \end{bmatrix}, \tag{S39}$$

where all commas in subscripts denote partial derivatives (i.e., $f_{,x} \equiv \frac{\partial f}{\partial x}$). With $\Delta x_{NP} = x - x_{NP}$ and $\Delta y_{NP} = y - y_{NP}$ expressions for each partial derivative in Equation (S39) are given by:

$$u_{z,x}(x,y,z) = \begin{cases} 0 & \Delta r_{NP} \le 0 \quad \text{and} \quad \Delta Z_{NP} \ge 0 \\ -2\frac{\Delta Z_{NP} \Delta r_{NP} \Delta x_{NP}}{r_{local}\sigma^2} \exp\left(-\frac{\Delta r_{NP}^2}{\sigma^2}\right) & \Delta r_{NP} > 0 \quad \text{and} \quad \Delta Z_{NP} \ge 0 \\ 0 & \Delta Z_{NP} < 0 \end{cases}, \tag{S40}$$

$$u_{z,y}(x,y,z) = \begin{cases} 0 & \Delta r_{NP} \le 0 \quad \text{and} \quad \Delta Z_{NP} \ge 0 \\ -2\frac{\Delta Z_{NP} \Delta r_{NP} \Delta y_{NP}}{r_{local}\sigma^2} \exp\left(-\frac{\Delta r_{NP}^2}{\sigma^2}\right) & \Delta r_{NP} > 0 \quad \text{and} \quad \Delta Z_{NP} \ge 0 \\ 0 & \Delta Z_{NP} < 0 \end{cases}, \tag{S41}$$

$$u_{z,Z}(x,y,z) = \begin{cases} -1 & \Delta r_{NP} \le 0 \quad \text{and} \quad \Delta Z_{NP} \ge 0 \\ -\exp\left(-\frac{\Delta r_{NP}^2}{\sigma^2}\right) & \Delta r_{NP} > 0 \quad \text{and} \quad \Delta Z_{NP} \ge 0 \\ 0 & \Delta Z_{NP} < 0 \end{cases}. \tag{S42}$$

The area stretch ratio (termed $\alpha$ in the main text) is given by the kinematic relationship [11]:

$$\alpha = \frac{A}{A_0} = \det(\mathbf{F}) \left| \mathbf{n} \cdot \mathbf{F}^{-1} \right| \tag{S43}$$

$$= (1 + u_{z,Z}) \left| \begin{bmatrix} n_x \\ n_y \\ n_z \end{bmatrix} \cdot \begin{bmatrix} 1 & 0 & 0 \\ 0 & 1 & 0 \\ -\frac{u_{z,x}}{1+u_{z,Z}} & -\frac{u_{z,y}}{1+u_{z,Z}} & \frac{1}{1+u_{z,z}} \end{bmatrix} \right| \tag{S44}$$

$$= (1 + u_{z,Z}) \sqrt{\left( n_x - \frac{n_z u_{z,x}}{1 + u_{z,Z}} \right)^2 + \left( n_y - \frac{n_z u_{z,y}}{1 + u_{z,Z}} \right)^2 + \left( \frac{n_z}{1 + u_{z,Z}} \right)^2}, \tag{S45}$$

where $\mathbf{n} = [n_x, n_y, n_z]$ is the normal vector of the reference surface:

$$\mathbf{n} = \frac{a_{nuc}^2 b_{nuc}^2}{\sqrt{(x^2 + y^2) b_{nuc}^4 + (Z - z_{nuc})^2 a_{nuc}^4}} \left[ \frac{x}{a_{nuc}^2}, \frac{y}{a_{nuc}^2}, \frac{Z - z_{nuc}}{b_{nuc}^2} \right]. \tag{S46}$$

Finally, substituting the above expressions yields:

$$\alpha = \begin{cases} n_z & \Delta r_{NP} \leq 0,\ \Delta Z_{NP} \geq 0 \\ (1 - \sigma_1) \sqrt{\left( n_x + \frac{2 n_z \Delta x_{NP} \sigma_1 \sigma_2}{1 - \sigma_1} \right)^2 + \left( n_y + \frac{2 n_z \Delta y_{NP} \sigma_1 \sigma_2}{1 - \sigma_1} \right)^2 + \left( \frac{n_z}{1 - \sigma_1} \right)^2} & \Delta r_{NP} > 0,\ \Delta Z_{NP} \geq 0\ , \\ 1 & \Delta Z_{NP} < 0 \end{cases} \tag{S47}$$

$$\text{where } \sigma_1 = \exp\left( -\frac{\Delta r_{NP}^2}{\sigma^2} \right) \quad \text{and} \quad \sigma_2 = \frac{\Delta Z_{NP} \Delta r_{NP}}{r_{local} \sigma^2}.$$

The curvature of the membrane directly above a nanopillar is identically zero. To compute the curvature of the nuclear membrane elsewhere when $\Delta Z_{NP} \geq 0$, we represent it as a Monge patch [12]:

$$z(x, y) = Z + \Delta Z_{NP} \exp\left( -\frac{\Delta r_{NP}^2}{\sigma^2} \right) \tag{S48}$$

For convenience, we first define the partial derivatives associated with the reference ellipsoidal surface:

$$\begin{aligned} Z_{,x} &= \frac{b_{nuc} x}{a_{nuc}^2 \alpha} \\ Z_{,y} &= \frac{b_{nuc} y}{a_{nuc}^2 \alpha} \\ Z_{,xy} &= \frac{b_{nuc} xy}{a_{nuc}^4 \alpha^3} \\ Z_{,xx} &= \frac{b_{nuc} x^2}{a_{nuc}^4 \alpha^3} + \frac{b_{nuc}}{a_{nuc}^2 \alpha} \\ Z_{,yy} &= \frac{b_{nuc} y^2}{a_{nuc}^4 \alpha^3} + \frac{b_{nuc}}{a_{nuc}^2 \alpha}, \\ \text{where} \quad \alpha &= \sqrt{1 - \frac{x^2 + y^2}{a_{nuc}^2}}. \end{aligned} \tag{S49}$$

The mean curvature is then given explicitly by:

$$H_{NE}(x,y) = -\frac{(1+z_{,y}^2)z_{,xx} - 2z_{,x}z_{,y}z_{,xy} + (1+z_{,x}^2)z_{,yy}}{2(1+z_{,x}^2+z_{,y}^2)^{3/2}}, \tag{S50}$$

where

$$\begin{aligned}
z_{,x} &= Z_{,x}(1-\sigma_1) - 2\sigma_1\sigma_2\Delta x_{NP} \\
z_{,y} &= Z_{,y}(1-\sigma_1) - 2\sigma_1\sigma_2\Delta y_{NP} \\
z_{,xy} &= Z_{,xy}(1-\sigma_1) + 2\sigma_1\sigma_2\Delta x_{NP}\Delta y_{NP}\left(\frac{Z_{,x}}{\Delta Z_{NP}\Delta x_{NP}} + \frac{Z_{,y}}{\Delta Z_{NP}\Delta y_{NP}} + \frac{2\Delta r_{NP}}{\sigma^2 r_{local}} + \frac{1}{r_{local}^2} - \frac{1}{r_{local}\Delta r_{NP}}\right) \\
z_{,xx} &= Z_{,xx}(1-\sigma_1) + 2\sigma_1\sigma_2\Delta x_{NP}^2\left(\frac{2Z_{,x}}{\Delta Z_{NP}\Delta x_{NP}} + \frac{2\Delta r_{NP}}{\sigma^2 r_{local}} + \frac{1}{r_{local}^2} - \frac{1}{r_{local}\Delta r_{NP}} - \frac{1}{\Delta x_{NP}^2}\right) \\
z_{,yy} &= Z_{,yy}(1-\sigma_1) + 2\sigma_1\sigma_2\Delta y_{NP}^2\left(\frac{Z_{,y}}{\Delta Z_{NP}\Delta y_{NP}} + \frac{2\Delta r_{NP}}{\sigma^2 r_{local}} + \frac{1}{r_{local}^2} - \frac{1}{r_{local}\Delta r_{NP}} - \frac{1}{\Delta y_{NP}^2}\right).
\end{aligned}$$

The above equations consider deformation ascribed to a single nanopillar acting on the nuclear envelope. To ensure continuity of membrane deformations in response to multiple nanopillars, we simply consider the summation of effects at a given point, provided it is not directly above a nanopillar:

$$u_z \leftarrow \sum_{n=1}^{N} \begin{cases} \Delta z_{NP} \exp\left(-\frac{(r_{local,n} - (r_{NP} + 2d_{steric}))^2}{\sigma^2}\right) & \Delta Z_{NP} \geq 0 \\ 0 & \Delta Z_{NP} < 0 \end{cases} \tag{S51}$$

where $N$ is the total number of nanopillars under consideration and $r_{local,n} = \sqrt{(x - x_{NP,n})^2 + (y - y_{NP,n})^2}$ is the distance from nanopillar $n$. Generally speaking, because $\sigma = 0.2$ µm, which is less than the spacing between nanopillars, the deformations are mainly attributed to the nanopillar closest to a given point on the nuclear envelope. If the point under consideration is in the upper region of the nuclear envelope (i.e., $Z_{NP} < 0$), then the curvature is given by setting $\sigma_1 = \sigma_2 = 0$ in Equation (S50).

To ensure conservation of volume for the nucleus, we adjust $a_{nuc}$ in Equation (S37) such that the deviation in volume is less than 1% after applying the indentation described above.

### Supporting Figures and Movies

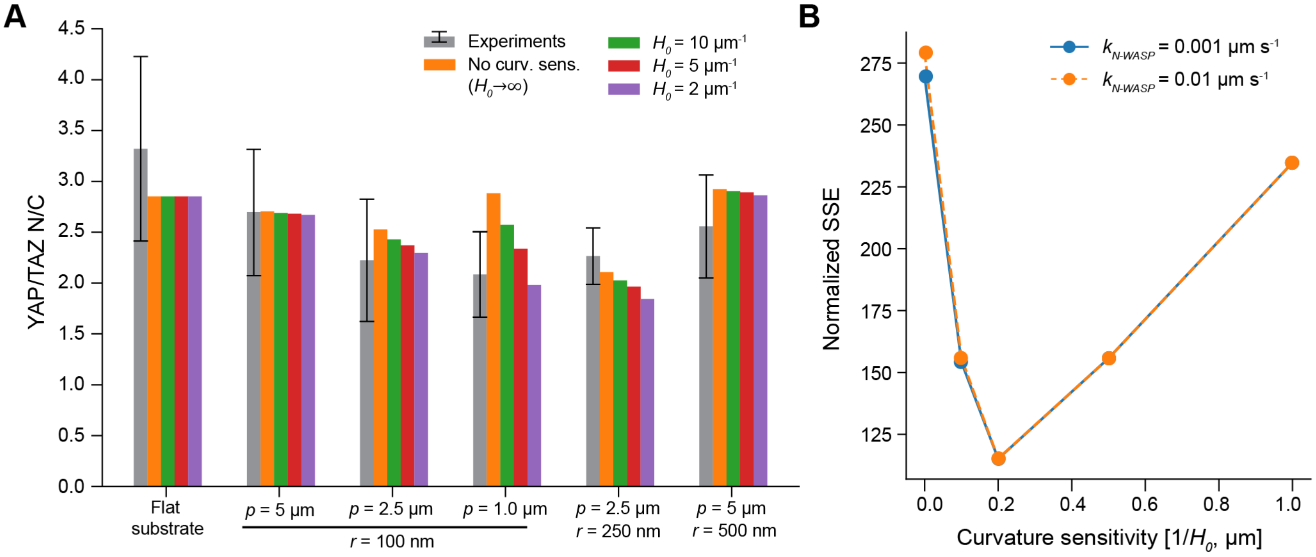


**Figure S1: Determination of best-fit FAK curvature sensitivity.** A) YAP/TAZ N/C in experiments (grey bars: mean ± standard deviation) and in simulations (spatially averaged at steady-state) with different levels of curvature sensitivity. B) Normalized SSE (Eq S9) as a function of curvature sensitivity ($1/H_0$) for two different rates of N-WASP-mediated actin assembly at regions of high PM curvature.

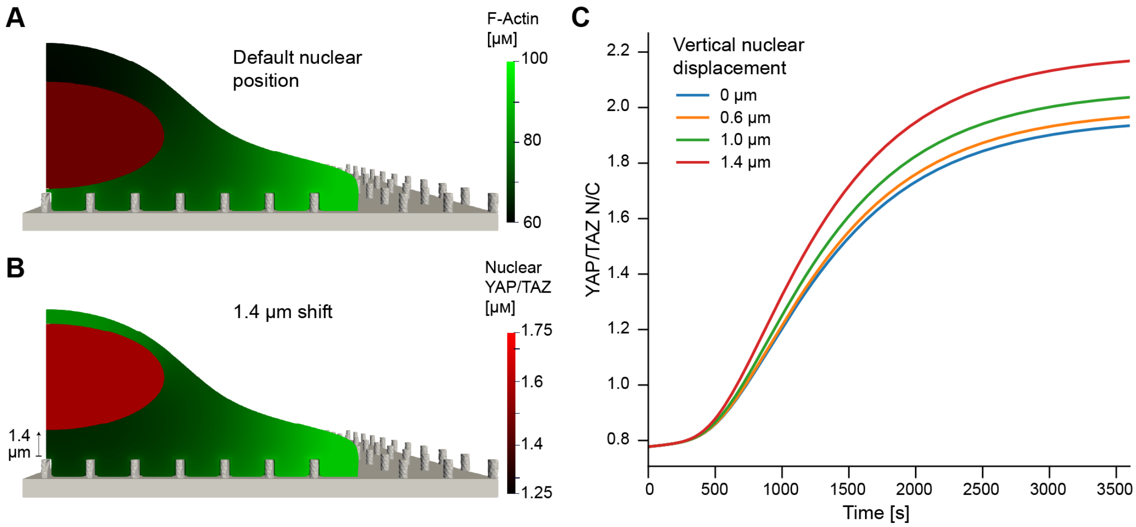


**Figure S2: Decreasing the gap between the nucleus and upper plasma membrane enhances YAP/TAZ nuclear entry.** A-B) Contrast in nuclear YAP/TAZ and F-actin distributions between the default nuclear position (A) and a nucleus shifted up by 1.4 μm (B) at steady-state ($t$ = 10,000 s). C) YAP/TAZ N/C dynamics for different nuclear locations, showing increased nuclear accumulation for larger vertical shifts. All simulations in this figure were conducted for a substrate with $r_{NP}$ = 250 nm, $p_{NP}$ = 2.5 μm, and $h_{NP}$ = 1 μm.

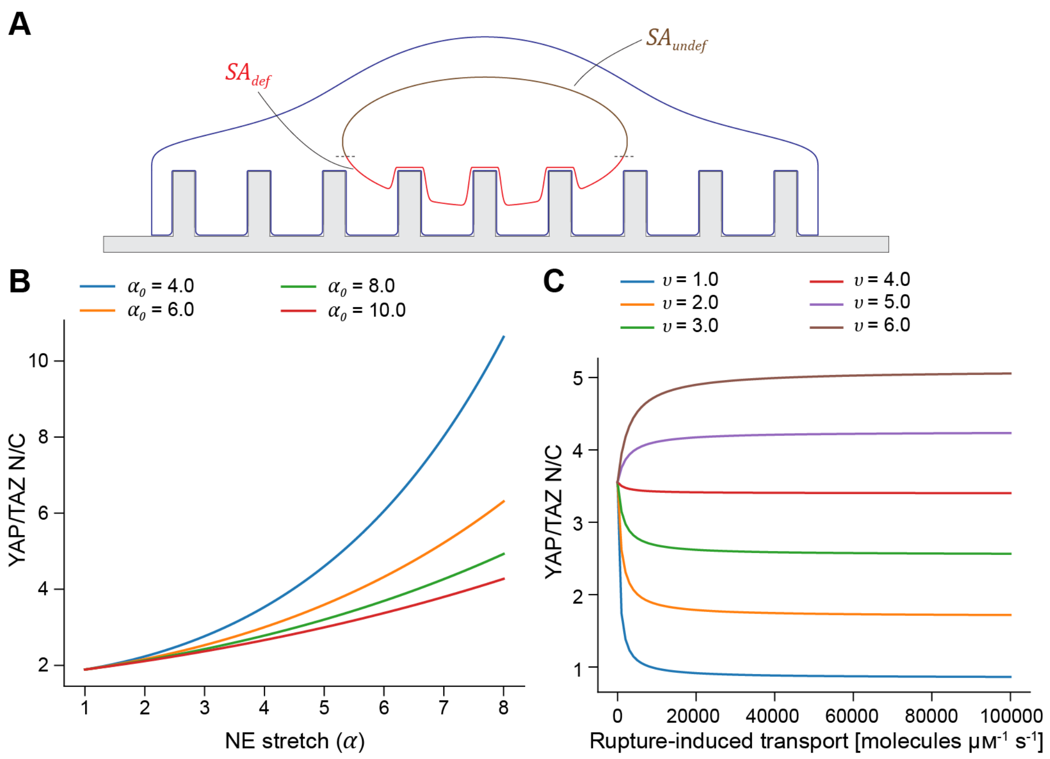


**Figure S3: Well-mixed model predictions for YAP/TAZ N/C.** A) Schematic showing two regions of the nuclear envelope (deformed and undeformed) considered in the well-mixed model. B) Predicted stretch-dependent effects on YAP/TAZ transport through NPCs using Eq 5 and $SA_{def} \approx \alpha SA_0$. Based on values in spatial simulations of indented nuclei, we use $\phi_{free} = 0.85$, $[NPC_A]_{def} = 0.3$, $[NPC_A]_{undef} = 0.1$, $SA_0 = 5$ μm$^2$, and $SA_{undef} = 40$ μm$^2$. C) Predicted NER-dependent effects on YAP/TAZ transport through NPCs using Eq 5. We use the same parameters as in panel B, but with $\alpha = 4.5$ and $\alpha_0 = 5$; the $x$-axis corresponds to $SA_{NER} k_{rupture}$.

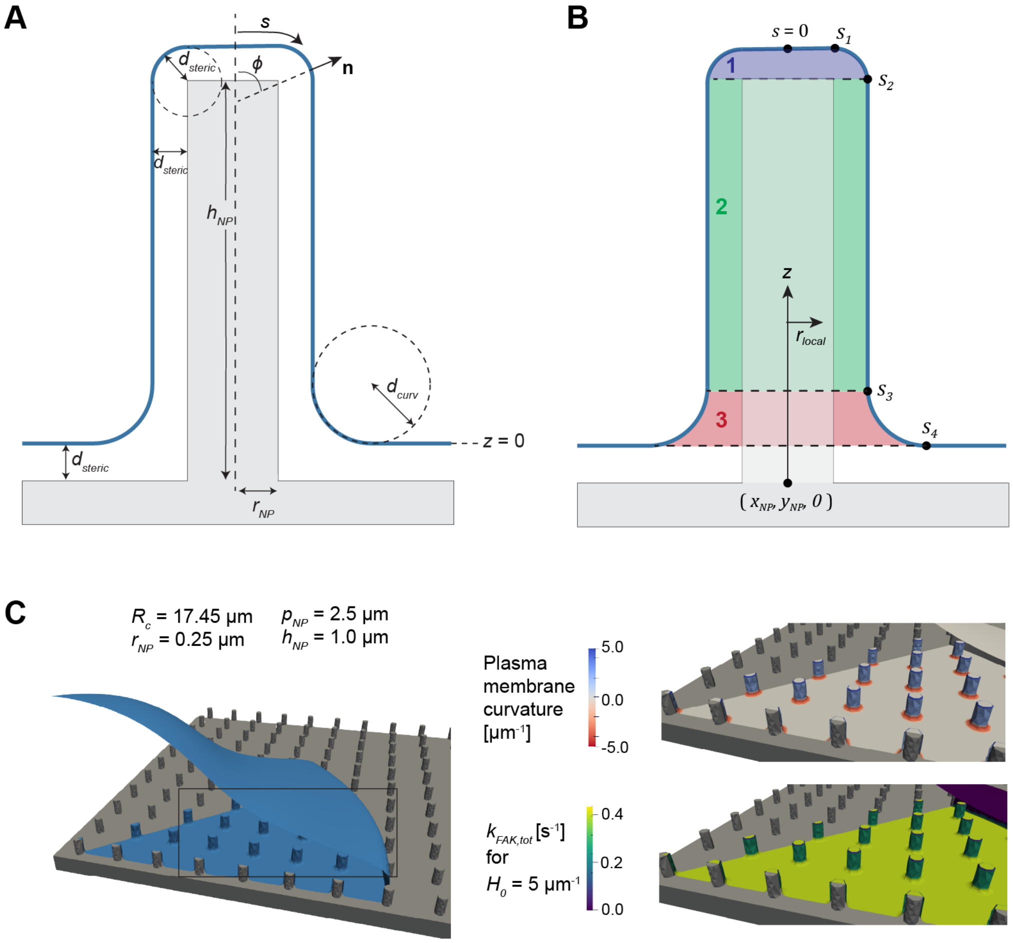


**Figure S4: Schematics of plasma membrane deformation on nanopillar substrates.** A) Cross-section of the PM (blue) near the surface of a cylindrical nanopillar. Relevant parameters required for calculation of the PM curvature (Eq S36) are defined graphically. This section of the geometry is axisymmetric with respect to the central axis of the nanopillar (dashed line). B) PM cross section with volumetric regions 1, 2, and 3 corresponding to those in Equation (S4) and arc length coordinates ($s_1$, $s_2$, $s_3$, and $s_4$) corresponding to those in Equations (S32) to (S36). C) A sample PM geometry (left, blue surface) on a substrate with 250 nm radius nanopillars ($p_{NP} = 2.5$ μm, $h_{NP} = 1.0$ μm), along with the resulting PM curvature (top right) and the local rate of curvature-dependent FAK phosphorylation when $H_0 = 5\text{μm}^{-1}$ (lower right).

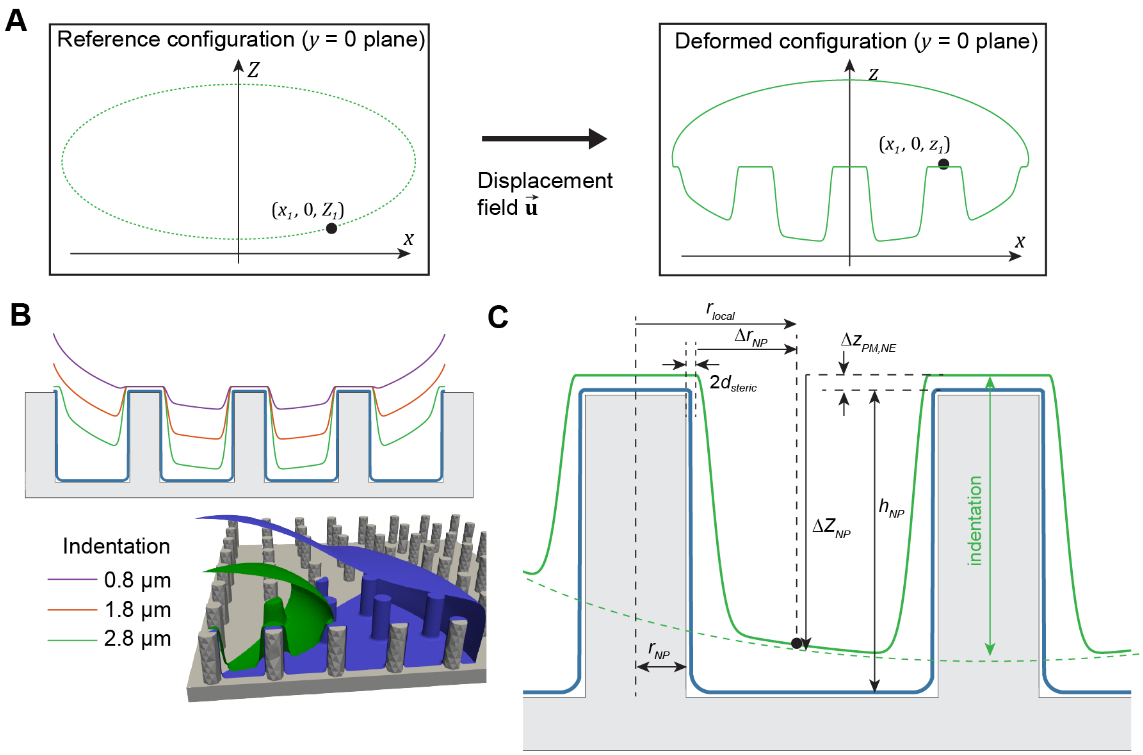


**Figure S5: Schematics of nuclear deformation on nanopillar substrates.** A) Summary of reference coordinates $(x, y, Z)$ (upper) vs. deformed coordinates $(x, y, z)$ (lower) over the $y = 0$ cross section of the nuclear envelope. B) A cross-section ($y = 0$) through the center of the cell showing prescribed deformations of the NE for different values of nuclear indentation (upper), with the 3D surface mesh associated with a deformation of 2.8 μm (lower). PM configuration is plotted in blue. C) Magnified schematic with local coordinates ($r_{local}$, $\Delta r_{NP}$, and $\Delta Z_{NP}$) defined with respect to the left nanopillar. $r_{local} = \sqrt{(x - x_{NP})^2 + (y - y_{NP})^2}$, $\Delta Z_{NP} = h_{NP} + \Delta z_{PM,NE} - Z$, and $\Delta r_{NP} = r_{local} - (r_{NP} + 2d_{steric})$.

**Movie 1:** Fast time-scale dynamics of F-actin in a cell on a substrate with 500 nm radius nanopillars (5.0 μm spacing, 1.0 μm height).

**Movie 2:** Fast time-scale dynamics of F-actin in a cell on a substrate with 250 nm radius nanopillars (2.5 μm spacing, 1.0 μm height).

**Movie 3:** Fast time-scale dynamics of F-actin in a cell on a substrate with 100 nm radius nanopillars (2.5 μm spacing, 1.0 μm height).

**Movie 4:** Spatial dynamics of cytosolic F-actin and nuclear YAP/TAZ in a cell on a flat substrate (substrate stiffness = 10 GPa).

**Movie 5:** Spatial dynamics of cytosolic F-actin and nuclear YAP/TAZ in a cell on a substrate with 100 nm radius nanopillars (2.5 μm spacing, 1.0 μm height).

**Movie 6:** Spatial dynamics of cytosolic F-actin and nuclear YAP/TAZ in a cell on a substrate with 100 nm radius nanopillars (5.0 μm spacing, 1.0 μm height).

**Movie 7:** Spatial dynamics of cytosolic F-actin and nuclear YAP/TAZ in a cell on a substrate with 500 nm radius nanopillars (3.5 μm spacing, 3.0 μm height) with no nuclear indentation.

**Movie 8:** Spatial dynamics of cytosolic F-actin and nuclear YAP/TAZ in a cell on a substrate with 500 nm radius nanopillars (3.5 μm spacing, 3.0 μm height) with 0.8 μm nuclear indentation.

**Movie 9:** Spatial dynamics of cytosolic F-actin and nuclear YAP/TAZ in a cell on a substrate with 500 nm radius nanopillars (3.5 μm spacing, 3.0 μm height) with 1.8 μm nuclear indentation.

**Movie 10:** Spatial dynamics of cytosolic F-actin and nuclear YAP/TAZ in a cell on a substrate with 500 nm radius nanopillars (3.5 μm spacing, 3.0 μm height) with 2.8 μm nuclear indentation.

**Movie 11:** Spatial dynamics of cytosolic and nuclear phosphorylated YAP/TAZ following nuclear envelope rupture (pore radius = 200 nm) in a cell with 2.8 μm nuclear indentation, assuming $\upsilon = 5$. The boundary of the nucleus is shaded to indicate NE permeability.

**Movie 12:** Spatial dynamics of cytosolic and nuclear phosphorylated YAP/TAZ following nuclear envelope rupture (pore radius = 200 nm) in a cell with 2.8 μm nuclear indentation, assuming $\upsilon = 1$. The boundary of the nucleus is shaded to indicate NE permeability.

12. Weisstein, E. W. *Monge Patch* https://mathworld.wolfram.com/MongePatch.html. Text.
